# A kinesin Klp10A mediates cell cycle-dependent shuttling of Piwi between nucleus and nuage

**DOI:** 10.1101/474049

**Authors:** Zsolt G. Venkei, Charlotte Choi, Suhua Feng, Steven E. Jacobsen, John K. Kim, Yukiko M. Yamashita

## Abstract

The piRNA pathway protects germline genomes through transcript cleavage of selfish genetic elements, such as transposons, in the cytoplasm and their transcriptional silencing in the nucleus. Here, we describe a mechanism by which the nuclear and cytoplasmic arms of the silencing mechanism are linked. During mitosis of *Drosophila* spermatogonia, nuclear Piwi interacts with nuage, the compartment that mediates the cytoplasmic arm of piRNA-mediated silencing. At the end of mitosis, Piwi leaves nuage to return to the nucleus. We found that dissociation of Piwi from nuage occurs at the depolymerizing microtubules of the central spindle, mediated by a microtubule-depolymerizing kinesin Klp10A. Depletion of *klp10A* delays Piwi’s return to the nucleus and affects piRNA production, suggesting the importance of nuclear-cytoplasmic communication in piRNA biogenesis. We propose that cell cycle-dependent communication between the nuclear and cytoplasmic arms of the piRNA pathway plays important roles in coordinated piRNA production.

## Introduction

Piwi-associated RNAs (piRNAs), a class of endogenous small RNAs found across multiple organisms, associate with Piwi proteins of the Argonaute family to silence active transposable elements (TEs)(Aravin et al., 2003; Batista et al., 2008; Brennecke et al., 2007; Girard et al., 2006; Ruby et al., 2006). In *Drosophila melanogaster*, Piwi, Aubergine (Aub), and Argonaute 3 (Ago3) comprise the three Piwi proteins of the piRNA pathway (Brennecke et al., 2007; Gunawardane et al., 2007; Nishida et al., 2007; Wang et al., 2015). The piRNA pathway is initiated in the nucleus, where primary piRNA precursors are transcribed from genomic piRNA clusters, processed, and translocated to the cytoplasm (Brennecke et al., 2007; Vagin et al., 2006). Once there, primary piRNAs are amplified in a “ping-pong” cycle that serves to simultaneously produce secondary piRNAs and cleave TEs (Gunawardane et al., 2007; Li et al., 2009; Wang et al., 2014). Specifically, primary piRNAs antisense to TEs are loaded into and direct Aub to mediate the cleavage of TEs, which produces secondary piRNAs that are subsequently loaded into Ago3. The secondary piRNAs generated by the ping-pong cycle are then loaded into Piwi and translocated into the nucleus as a piRNA-induced silencing complex (piRISC) (Senti et al., 2015; Wang et al., 2015). Within the nucleus, Piwi leads to the transcriptional repression of TEs through the deposition of heterochromatic histone marks (Klenov et al., 2014; Le Thomas et al., 2013; Sienski et al., 2015). Thus, the cytoplasmic arm of the piRNA pathway intimately interacts with the nuclear arm to affect silencing at the post-transcriptional and transcriptional levels, respectively.

As evidence of the tight coupling between the two arms of the piRNA pathway, ping-pong amplification takes place in nuage, electron-dense, phase-separated granules anchored directly to the cytoplasmic face of the nuclear pore (Lim and Kai, 2007; Malone et al., 2009; Zhang et al., 2012). The interface between nuage and the nucleus has been shown to be critical for piRNA biogenesis and transposon target recognition and repression. Although nuage has been characterized as a static and long-term platform for piRNA biogenesis, individual nuage components, such as Aub and Ago3, are dynamically shuttled in and out of these granules (Andress et al., 2016; Brennecke et al., 2007; Ryazansky et al., 2016; Snee and Macdonald, 2004; Wang et al., 2015). Unlike Aub and Ago3, Piwi shows predominantly nuclear localization (Gonzalez et al., 2015) and has not been well-characterized within the context of nuage. Although previous studies have identified multiple piRNA pathway components that are required for proper localization of piRNA amplification machinery to nuage granules, including Spindle-E, Qin, Krimper, and Vasa, the underlying mechanism by which ping-pong-amplified piRNAs translocate into the nucleus remains poorly understood (Andress et al., 2016; Sato et al., 2015; Wang et al., 2015; Webster et al., 2015; Zhang et al., 2012; Zhang et al., 2011).

Here we show that nuclear and cytoplasmic components of the piRNA pathway interact specifically during mitosis of *Drosophila* spermatogonia (SGs) to facilitate communication between the nuclear and cytoplasmic arms of the pathway. We found that Piwi colocalizes to nuage specifically during mitosis and then returns to the nucleus upon mitotic exit. Through immunoprecipitation and mass spectrometry, we found that Klp10A, a MT-depolymerizing kinesin motor protein (Rogers et al., 2004), physically interacts with many components of the piRNA pathway. In the absence of Klp10A, Piwi remains in nuage for a prolonged period after mitotic exit, thus delaying its return to the nucleus. Cytological observations suggest that dissociation of Piwi from nuage at the end of mitosis is facilitated at the central spindle while it is being depolymerized in a *klp10A*-dependent manner. We observed an upregulation of TE transcripts in early germ cells, likely due to a failure in Piwi-mediated transcriptional silencing. Increased TE transcript levels in early germ cells, combined with intact cytoplasmic ping-pong machinery, leads to an amplified ping-pong signature, which later leads to enhanced TE repression in spermatocytes (SCs). Taken together, we propose that interaction of Piwi with nuage during mitosis may allow Piwi to be loaded with an updated repertoire of piRNAs, targeting currently active TEs efficiently. We further propose that Klp10A plays a critical role in this process by facilitating dissociation of Piwi from nuage at the end of mitosis.

## Results

### Klp10A physically interacts with piRNA pathway components in germline stem cells and spermatogonia of *Drosophila* testis

In a previous study, we showed that Klp10A protein is enriched on the centrosomes specifically in germline stem cells (GSCs), but not in differentiating SGs, in the *Drosophila* testis (Chen et al., 2016). Klp10A protein also localizes to central spindle in both GSCs and SGs. Because centrosome play a critical role in asymmetric stem cell divisions through their unique behavior in stem cells of many model systems (Cheng et al., 2008; Conduit and Raff, 2010; Inaba et al., 2015b; Januschke et al., 2011; Januschke et al., 2013; Salzmann et al., 2014; Venkei and Yamashita, 2015; Wang et al., 2009; Yamashita et al., 2003; Yamashita et al., 2007), stem cell-specific centrosomal components are of significant interest. In an attempt to isolate proteins that specifically localize to GSC centrosomes, we affinity-purified Klp10A from a GSC-enriched extract: Klp10A was tagged with green fluorescent protein (GFP) and anti-GFP antibody was used to immunoprecipitate Klp10A and any associated proteins. To enrich for GSCs, self-renewal factor Unpaired (Upd) (Kiger et al., 2001; Tulina and Matunis, 2001) was co-expressed together with Klp10A-GFP (*nos-gal4>UAS-klp10A-GFP, UAS-upd*) (Figure 1A). Klp10A-GFP recapitulated the localization of endogenous Klp10A visualized by anti-Klp10A antibody, i.e. at the GSC centrosomes (Figure 1B, arrow) and the central spindle in telophase of GSCs and SGs (Figure 1B, arrowhead) (Chen et al., 2016).

**Figure 1.**
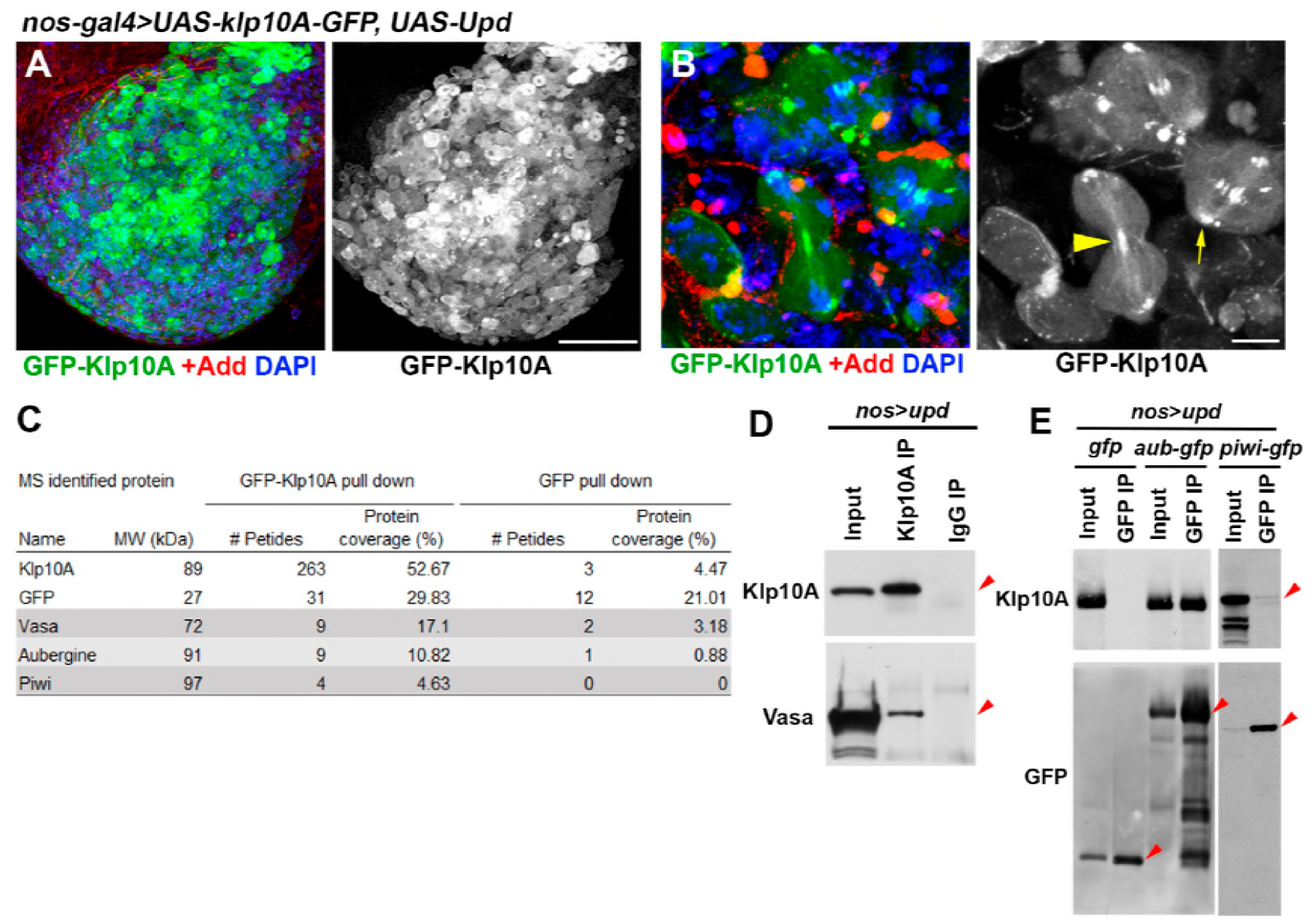
GFP-Klp10A localizes to the central spindle and interacts with piRNA pathway components in GSCs. A) Expression of GFP-Klp10A (green) in the *upd*-expressing testes (*nos-gal4>UAS-upd, UAS-gfp-klp10A*). Adducin (red), DAPI (blue). Bar: 50 *μ*m. B) GFP-Klp10A localization in *upd*-expressing germ cells. Arrow indicates localization at the centrosome, arrowhead indicates localization at the central spindle. Bar: 5 *μ*m. C) Summary of mass-spectrometry hits of piRNA pathway components from GFP-Klp10A and GFP co-IPs. D) co-immunoprecipitation of endogenous Vasa and Klp10A from *nos-gal4>UAS-upd* testes. E) co-immunoprecipitation of endogenous Klp10A with GFP-Aub or -Piwi in *upd-* expressing testes (*nos-gal4>UAS-upd*). D-E) Arrowheads point to the position of indicated proteins.

Anti-GFP immunoprecipitates were analyzed by mass-spectrometry (see Methods). To our surprise, we found that components of the piRNA pathway, such as Vasa, Aub and Piwi, were enriched in the Klp10A-GFP pull-down (Figure 1C, Figure 1-Source Data 1). We also found that many mRNA binding proteins were enriched (Figure 1-Source Data 1), details of which will be reported elsewhere. Immunoprecipitation using anti-Klp10A antibody from GSC-enriched extracts confirmed that Klp10A physically interacts with Vasa (Figure 1D). Similarly, anti-GFP antibody pull down of Aub-GFP or Piwi-GFP confirmed their interaction with Klp10A (Figure 1E).

The Klp10A interaction with piRNA pathway components is not unique to GSCs, but was also maintained in SGs at the central spindle during telophase (see below). Thus, the study diverged from our initial intention to isolate GSC centrosome-specific components. However, we investigated the unexpected role of *klp10A* in the regulation of the piRNA pathway, as the study revealed an unappreciated mode of regulation of piRNA biogenesis.

### Depletion of Klp10A results in alteration of piRNA biogenesis

To explore whether Klp10A interaction with piRNA pathway components was functionally significant, we examined whether *klp10A* is required for repression of TEs or piRNA biogenesis (Toth et al., 2016). To characterize the role of *klp10A* in the piRNA pathway, we first performed deep sequencing of small RNAs from wild-type and *klp10A*-knockdown germ cells (*nos-gal4>UAS-klp10A^RNAi^*, validated in our previous study (Chen et al., 2016), referred to as *klp10A^RNAi^*) (Figure 2, Figure supplement 3). We defined piRNAs as reads of 23-29 nucleotides (nt) in length that did not map to microRNAs or ribosomal RNAs. To assess the global changes in piRNA expression upon *klp10A* loss, we profiled piRNA reads across the *Drosophila* transcriptome. The knockdown of *klp10A* by RNAi caused specific upregulation of piRNAs mapping to piRNA clusters and repetitive elements, with minimal differences in other genomic classes (Figure 2A), suggesting that *klp10A* may play a specific role in the piRNA pathway. When we analyzed the abundance of piRNAs mapping to TEs upon loss of *klp10A*, we observed dramatic upregulation of piRNA reads both antisense and sense to TEs (Figure 2B). To probe the effect of *klp10A* loss further, we analyzed the distribution of piRNA abundance across the transposon transcript *HOBO.* Both antisense and sense reads across *HOBO* were significantly upregulated in the *klp10A^RNAi^* samples compared to wild-type without any noticeable change in their distribution (Figure 2C).

**Figure 2.**
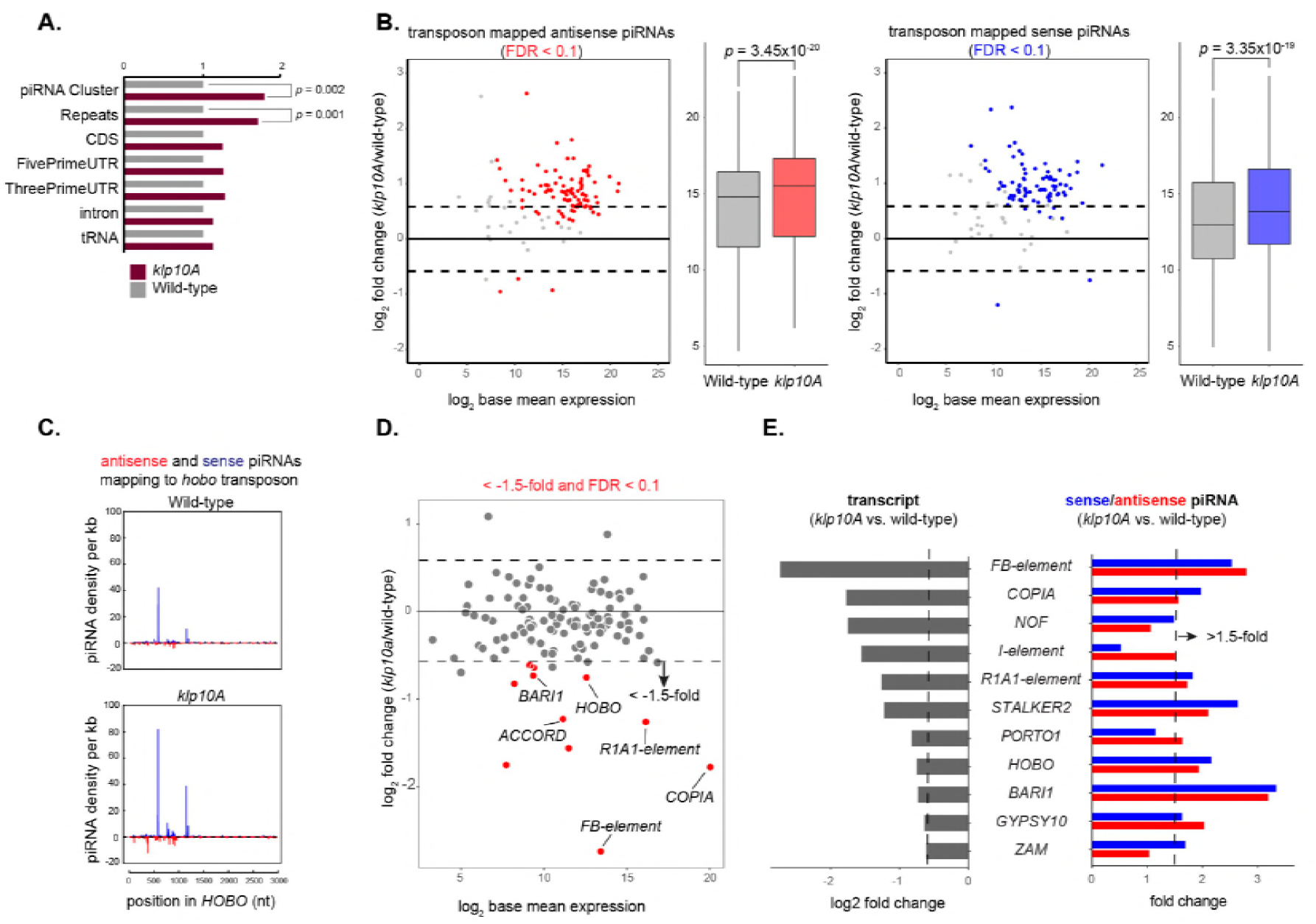
piRNA production is upregulated in *klp10A^RNAi^* testis. A) piRNA reads normalized to total library reads at each genomic feature. piRNAs are significantly upregulated in repeat and piRNA cluster regions. Significance was calculated using a student t-test. B) Scatter and bar plots comparing transposon mapped sense and antisense piRNA abundance in germ cells undergoing *klp10A^RNAi^* knockdown versus wild-type. A change of greater than 1.5-fold and adjusted p-value of 0.1 were used as cutoffs for differential analysis. p values for box plots were calculated using Wilcoxon signed-rank test. C) Density of sequenced piRNAs (blue: sense; red: antisense) across *HOBO*. The y-axis indicates relative units. Each piRNA read was normalized to miRNA hair-pin abundance. D) Scatter plot comparing transposon abundance in *klp10A^RNAi^* versus wild-type (red: FDR<0.1). E) Left bar plot show log2 fold change transcript abundance difference in *klp10A^RNAi^* versus wild-type of transposons with the greatest significant (FDR>0.1) downregulation in *klp10a* mutants versus wildtype. Right bar plot shows antisense and sense piRNA fold change differences in *klp10A^RNAi^* compared to wildtype at those transposons.

The above data showed that both primary and secondary piRNAs were upregulated. To test whether the ping-pong cycle was affected in *klp10A^RNAi^*, we calculated the number of antisense and sense read pairs that have complementary base pairing from their 5’ ends in both wildtype and *klp10A^RNAi^* libraries (Figure 2C, Figure 2-Figure Supplement 1A). We observed a significant bias for piRNA read pairs with 10-nucleotide complementarity in both wildtype and *klp10A^RNAi^*, suggesting that the ping-pong pathway remains active upon loss of *klp10A*. Moreover, we found no significant changes in ping-pong ratios in *klp10A^RNAi^* compared to wildtype (Figure 2 - Figure Supplement 1B), further confirming an intact ping-pong cycle. Given that loss of *klp10A* does not affect ping-pong amplification, the upregulation of both antisense and sense piRNAs seen in *klp10A^RNAi^* germ cells could be due to increased expression of piRNA precursors such as those generated from piRNA clusters and TEs.

To address if *klp10A^RNAi^* leads to changes in TE expression, we performed mRNA-sequencing in wildtype and *klp10A^RNAi^* germ cells (Figure 2-figure supplement 2). We found that loss of *klp10A* resulted in at least a 1.5-fold reduction in the levels of a subset of TEs, including *HOBO* (Figure 2D). In contrast, the majority of piRNAs that map to these TEs were upregulated (Figure 2E), consistent with the general trend of piRNA upregulation (Figure 2B). Based on these results, we speculate that moderate downregulation of TEs could be due to global upregulation of piRNAs in *klp10A^RNAi^*.

To further assess these results, we conducted single molecule fluorescent RNA *in situ* hybridization for *HOBO* transposon, an example of TEs whose transcription was downregulated but the corresponding piRNAs exhibited upregulation in *klp10A^RNAi^*. *HOBO* expression was quantified by counting the number of single molecule FISH signals. Interestingly, contrary to the prediction based on the sequencing analysis that showed downregulation of *HOBO* among other TEs, *HOBO* was transiently upregulated in SGs and early SCs (Figure 3A, B, D). This was followed by significant downregulation of *HOBO* in the SC stage (Figure 3A, B, D). This is in contrast to *aub^RNAi^*, which leads to upregulation of TEs through all stages of germ cell development (Figure 3C, E, Figure 3 - Figure3 supplement 1). Note that *HOBO* upregulation/downregulation at different stages of germ cell stages is not due to differential efficiency of *klp10A^RNAi^*, as Klp10A protein was undetectable in all stages of germ cells (Figure 3 - Figure3 supplement 2). Combined with the sequencing analysis that showed upregulation of the ping-pong signature and overall downregulation of transposon expression, we interpret these results to indicate that the TEs are slightly upregulated in *klp10A^RNAi^* SGs and early SCs, which leads to enhanced piRNA production through intact ping-pong cycle, which later downregulates the target mRNA in the late SCs (Figure 3F). Overall downregulation of TE expression revealed by the RNA sequencing of total testis is likely explained by the fact that there are more SCs than SGs in the testis.

**Figure 3.**
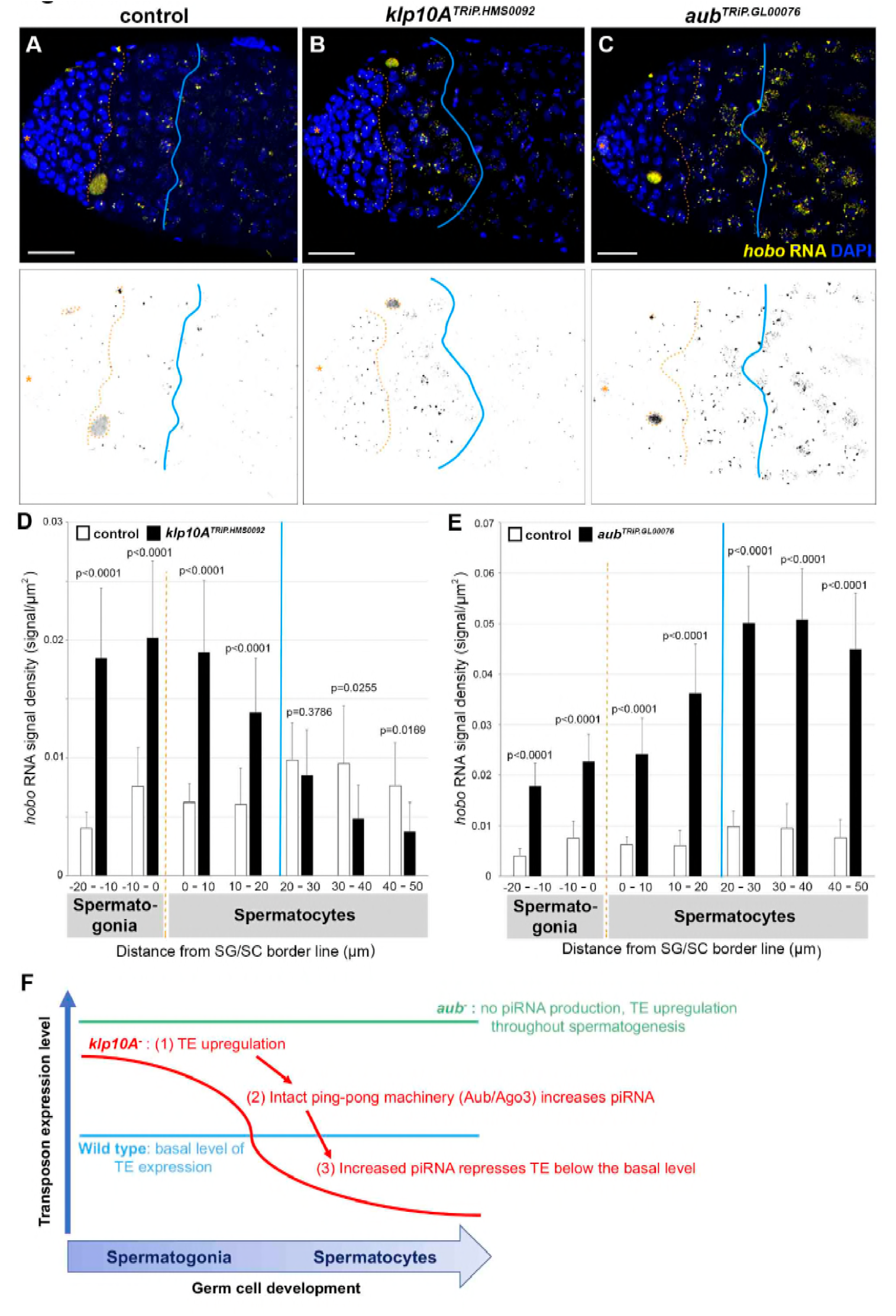
Transient upregulation and downregulation of *HOBO* upon *klp10A* knockdown. A-C) single molecule RNA *in situ* hybridization signals in wild type (A), *nos*>*klp10A^RNAi^* (B), and *nos*>*aub^RNAi^* testes (C). *HOBO* RNA (yellow), DAPI (blue). Asterisk indicates the hub. Orange dotted line indicates the border line between SG and SC cells. Blue line indicates the border between early SC and late SC, where *klp10A^RNAi^* transitions from up-to down-regulation of *HOBO* transcript. Dying cysts that take up *in situ* probe signals non-specifically (orange dotted circles) were manually removed from the analysis. Bars 20 μm. D-E) Density of *HOBO* RNA signals in SGs and early SCs after *klp10A^RNAi^* (D), or *aub^RNAi^* (E). *N*=9-9 testes for each genotype. Error bars indicate SD. p values of t-tests are provided. F) Model of *klp10A^RNAi^* effect on transposon expression in SGs and SCs.

Taken together, these results demonstrate that Klp10A’s interaction with piRNA pathway components indeed has functional significance, prompting us to further examine the underlying mechanisms. The above data specifically points to the role of *klp10A* in repression of TEs in SGs, without affecting the ping-pong cycle.

### Klp10A colocalizes with piRNA pathway components at the central spindle during telophase in GSCs and SGs

To begin to explore how *klp10A* might contribute to piRNA biogenesis, we examined the localization of Klp10A protein and piRNA pathway components throughout the cell cycle. Vasa and Aub showed well-established perinuclear localization in nuage throughout interphase (Figure 4A, B, Figure 4 – Figure supplement 1A, B) (Kibanov et al., 2011; Nagao et al., 2010; Nishida et al., 2007). During this period, Klp10A showed centrosome localization in GSCs (Figure 4A), or uniform cytoplasmic localization in SGs (Figure 4B) as we reported previously (Chen et al., 2016). We did not observe clear localization of Klp10A to nuage, suggesting that the potential role of Klp10A in piRNA pathway is not related to its GSC-specific centrosomal localization.

**Figure 4.**
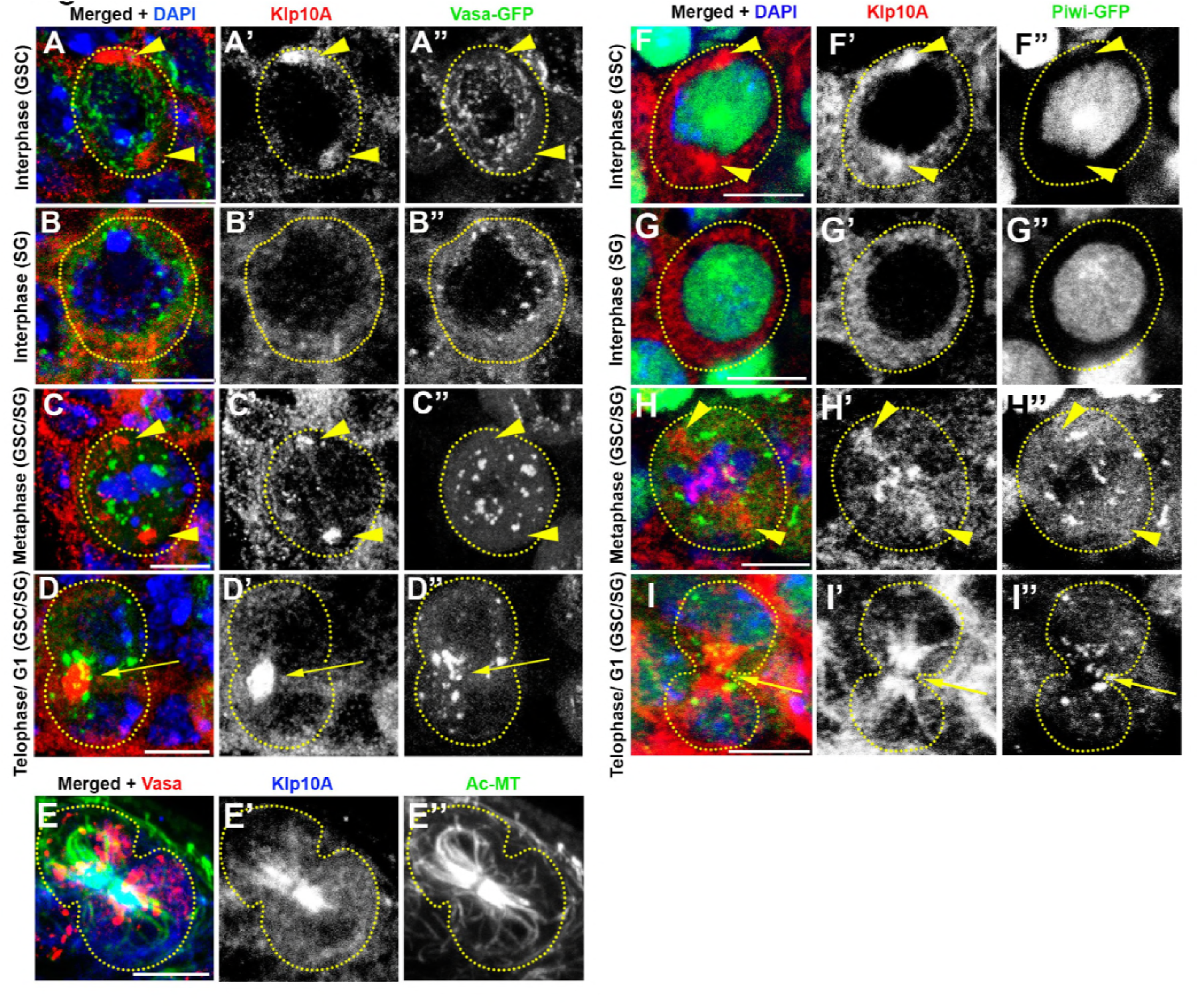
Vasa, Piwi and Klp10A colocalize at the central spindle of mitotic germ cells. A-D) Localization of Klp10A (red) and Vasa-GFP (green), counter-stained with DAPI (blue). Interphase GSC (A-A”), Interphase SG (B-B”), GSC/SG in metaphase (C-C”), and GSC/SG in telophase/G1(D-D”). E) Localization of Vasa (red), acetylated MTs (acMTs) (green) and Klp10A (blue) in a telophase GSC-GB (gonialblast) pair of a wild type testis. F-I) localization of Klp10A (red) and Piwi-GFP (green), counter-stained with DAPI (blue) in interphase GSC (F-F”), interphase SG (G-G”), metaphase GSC/SG (H-H”), and telophase/G1 GSC/SG (I-I”). Germ cells are indicated by dotted lines. Centrosomal localization of Klp10A is indicated by arrowheads, central spindle localization with arrows. Bars: 5 *μ*m.

When cells enter mitosis, nuage became somewhat larger in size, which we refer to as ‘mitotic nuage’ hereafter (Figure 4C, Figure 4 – Figure supplement 1C). Mitotic nuage appears to be the same population described previously as ‘Vasa granule near the mitotic chromosomes’ by Pek and Kai (Pek and Kai, 2011). At this point, Klp10A was observed on the spindle pole (and weakly on the spindle) as previously reported (Chen et al., 2016), showing little colocalization between Klp10A and Vasa/Aub (Figure 4C, Figure 4 – Figure supplement 1C).

Although Klp10A did not noticeably colocalize with any of nuage components until anaphase, colocalization became clear during telophase: nuage was always associated with Klp10A near the center of the dividing cell (Figure 4D, Figure 4 – Figure supplement 1D), which we confirmed to be bundles of central spindle MTs (Figure 4E). Importantly, Klp10A’s localization to the central spindle was observed both in GSCs and SGs (Chen et al., 2016), and colocalization of Klp10A and piRNA pathway components was commonly observed both in GSCs and SGs.

In contrast to Aub and Ago3, which reside in nuage together with Vasa to function in the ‘cytoplasmic arm’ of the piRNA pathway, Piwi functions in the nucleus to repress TEs at a transcriptional level (Aravin et al., 2008; Le Thomas et al., 2013; Nishida et al., 2015; Toth et al., 2016; Xiol et al., 2014; Zhang et al., 2012). We confirmed that Piwi-GFP localizes to the nucleus during interphase of GSCs and SGs as described previously (Cox et al., 2000; Gonzalez et al., 2015)(Figure 4F, G). Interestingly, we found that once cells enter mitosis, Piwi-GFP localized to mitotic nuage together with the components of the cytoplasmic arm of the piRNA pathway, such as Aub and Ago3 (Figure 4H, I, see Figure 5 for the confirmation of Piwi’s localization to mitotic nuage). During telophase, Piwi-GFP still colocalized with nuage components at the central spindle, after which it returned to the nucleus. Taken together, these data show that piRNA pathway components likely interact with Klp10A at the central spindle during telophase.

**Figure 5.**
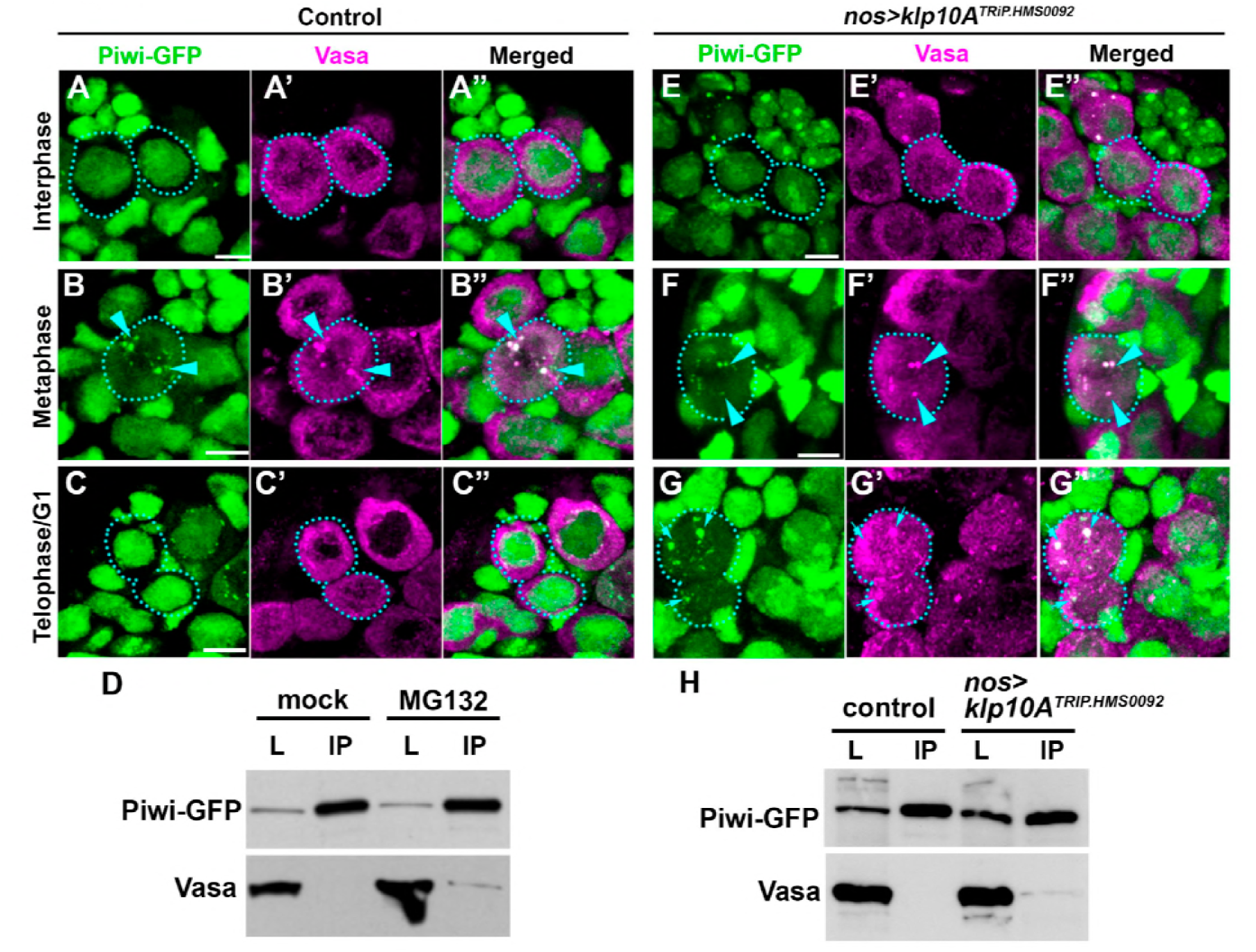
Piwi interacts with cytoplasmic nuage components in mitosis, and returns to the nucleus at the end of mitosis in a Klp10A-dependent manner. (A-C) Piwi-GFP (green) and Vasa (red) localization throughout the GSC cell cycle in wild type testes. The localization patterns were the same in GSCs and SGs. GSCs are encircled by dotted lines. Piwi-GFP and Vasa colocalization at the mitotic nuage is indicated by arrowheads. Bars: 5 *μ*m. D) co-immunoprecipitation of Vasa with Piwi-GFP from *nos>upd* testes after 4h 30min *ex vivo* MG132 treatment. (E-G) Piwi-GFP (green) and Vasa (red) localization throughout the GSC cell cycle in *klp10A^RNAi^* testes. Cytoplasmic Piwi-GFP/Vasa granule in post-mitotic interphase cells are highlighted by arrows in panel G. H) co-immunoprecipitation of Vasa with Piwi-GFP from *nos>upd* testes after *klp10A^RNAi^*.

### Klp10A is required for Piwi’s relocation from mitotic nuage to the nucleus at the end of mitosis

Having established that Klp10A interacts with nuage components at the central spindle, we explored how this interaction may impact the function of piRNA pathway, explaining enhanced ping-pong signature in *klp10A^RNAi^* testis. To this end, we first examined the localization of Piwi and Vasa in control and *klp10A^RNAi^* GSCs/SGs.

We confirmed our above observation that Piwi localizes to mitotic nuage by examining the colocalization of Piwi-GFP with Vasa. Although their localization was distinct during interphase (i.e. Vasa in nuage, Piwi in the nucleus) (Figure 5A), Vasa and Piwi colocalized during mitosis (Figure 5B), and Piwi returned to the nucleus at the end of mitosis (Figure 5C). We further detected physical interaction between Vasa and Piwi specifically when cells were arrested in mitosis by the use of MG132 (a proteasome inhibitor that arrests cells in mitosis (Moutinho-Pereira et al., 2010)) (Figure 5D). These results establish that Piwi gains access to nuage specifically during mitosis.

Next, we investigated whether *klp10A^RNAi^* testes show any defects in the behavior of nuage or Piwi during cell cycle. Nuage morphology and composition appeared unperturbed in most of interphase through anaphase in both GSCs and SGs in *klp10A^RNAi^* testes (Figure 5E, F). However, the difference between the control and *klp10A^RNAi^* germ cells became clear during telophase. In control germ cells, Piwi returned to the nucleus at the end of telophase and became exclusively nuclear in the subsequent interphase as described above (Figure 5C). In contrast, in *klp10A^RNAi^* germ cells, Piwi persisted in cytoplasmic nuage even after the completion of mitosis (Figure 5G). Piwi-Vasa interaction was detectable by co-immunoprecipitation in *klp10A^RNAi^* without being arrested in mitosis (Figure 5H), consistent with persistent interaction of Piwi with Vasa at nuage in interphase cells. Likely reflecting the delayed/incomplete translocation of Piwi from mitotic nuage to the nucleus, we also observed that the amount of nuclear Piwi was slightly but significantly reduced in GSCs/SGs in *klp10A^RNAi^* germ cells (Figure 5 – Figure Supplement 1). Delayed dissociation of Piwi from mitotic nuage at the end of mitosis was confirmed by live observation of Piwi-GFP (Movie 1 and 2).

Taken together, these results show that 1) Piwi interacts with nuage components (Vasa, Aub) specifically in mitosis, and 2) *klp10A* is required for releasing Piwi from mitotic nuage to allow its return to the nucleus at the end of mitosis.

### Piwi dissociates from nuage at the central spindle microtubules

How does Piwi dissociate from mitotic nuage at the end of mitosis, and how does *klp10A* promote this process? A closer examination of live-imaging observation using Piwi-GFP and Vasa-mCherry (Movie1) revealed that a Piwi-positive compartment (‘Piwi granule’ hereafter) budded off from mitotic nuage, then dispersed near the center of the telophase cell at the central spindle (see below for the confirmation of their localization at the central spindle) (Figure 6A, Movie 1). Concomitantly, nuclear Piwi level gradually increased (Movie1), suggesting that Piwi released from mitotic nuage returned to the nucleus. Morphology of nuage marked by Vasa remained unchanged during this period (Movie1). In contrast to control SGs, we barely observed the budding off of the Piwi granule in *klp10A^RNAi^* SGs (Figure 6B, Movie 2). As a result, Piwi stayed with nuage for a prolonged time period even after completion of mitosis. These results show that *klp10A* is required for releasing Piwi from mitotic nuage such that Piwi can return to the nucleus.

**Figure 6.**
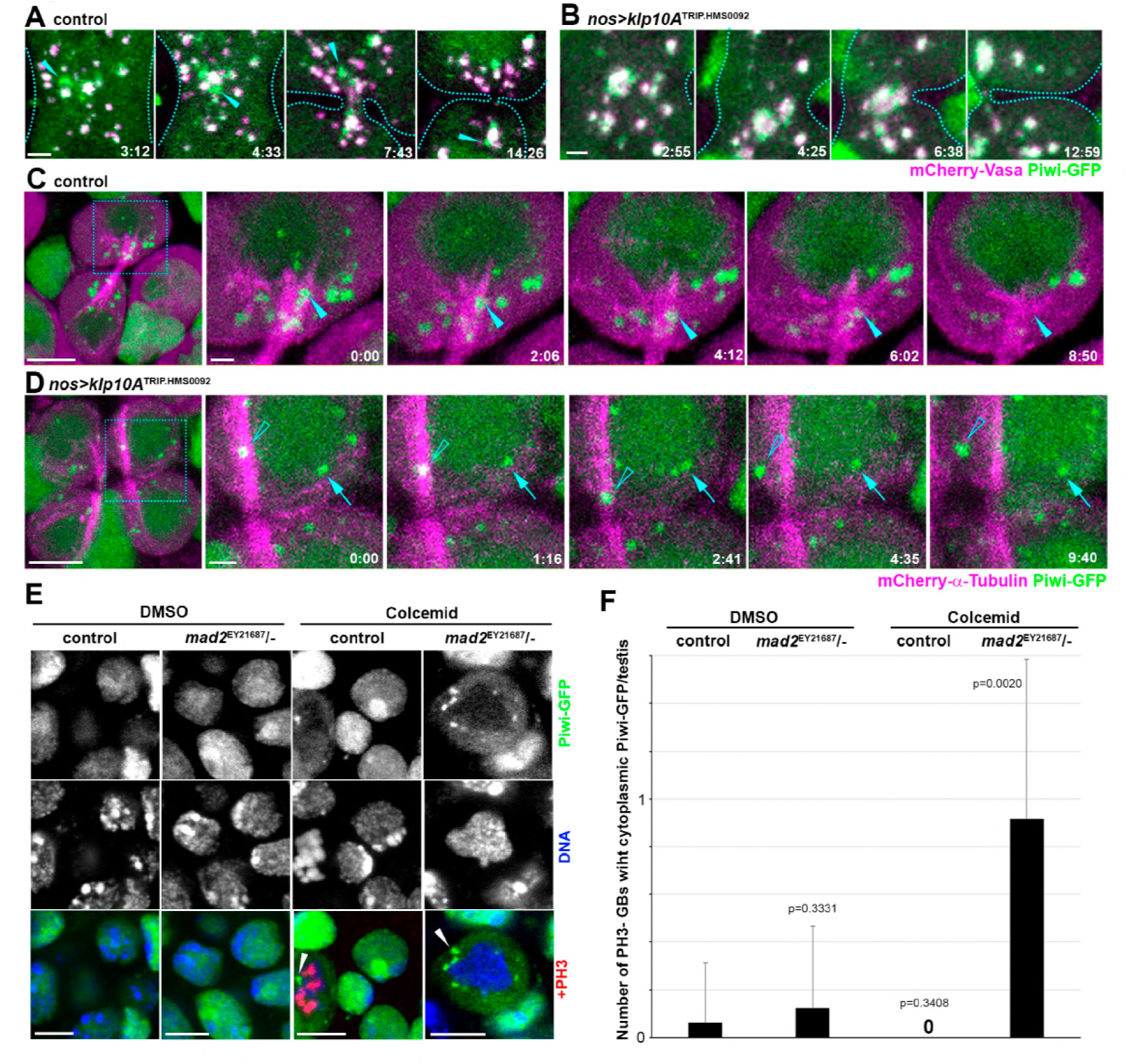
Dissociation of Piwi from mitotic nuage occurs at the central spindle in a Klp10A-dependent manner. A-B) Live imaging of mCherry-Vasa (red) and Piwi-GFP (green) in the central region of interconnected daughter SGs in wild type (A) and in *klp10A^RNAi^* (B) testes. The elapsed time after anaphase-B onset is indicted in min:sec. arrowheads point to Piwi-GFP signals budding off from the mitotic nuage. Dotted lines indicate the outline of cells. Bars: 1 *μ*m. C-D) Live imaging of mCherry-α-Tubulin (red) and Piwi-GFP (green) in post-mitotic SG cells of wild type (C) and *klp10A^RNAi^* (D) testes. Squares in the first panels indicate the zoomed regions shown by the later panels. Arrowheads in (C) point to a mitotic nuage particle, initially positive for Piwi and gradually losing the signal. Open arrowheads (D) point to a Piwi positive nuage particle, sliding along and later releasing the MT bundle. Bars: 5 *μ*m, and 1 *μ*m for zoomed regions. (E) Piwi-GFP (green) localization in interphase SGs after 3h *ex vivo* mock or colcemid treatment of wild type and *mad2* null mutant spermatogonia. Arrowheads point to Piwi-GFP positive nuage. DAPI (blue), PH3 marks mitotic chromatin (red). (F) Frequency of interphase (PH3-) GBs with Piwi-GFP positive nuage 3h after *ex vivo* colcemid treatment. Error bars indicate SD, *n*=13-18 testes per each of the genotype-treatment combination categories. p values of t-tests are provided.

The above live-imaging observation indicated that budding off of the Piwi granule from mitotic nuage occurs near the center of dividing cells (Movie1). Because Klp10A localizes to the central spindle and nuage components likely interact with Klp10A at the central spindle (Figure 4), we next examined how budding of the Piwi granule occurs at the central spindle. Using Piwi-GFP combined with mCherry-α-Tubulin, we found that the Piwi-GFP signal was associated with the central spindle MTs and moved back and forth along MTs, gradually decreasing in intensity until it completely dissipated (Figure 6C and Movie 3). In contrast, in *klp10A^RNAi^* GSCs/SGs, Piwi granule failed to disappear as they move along the central spindle MTs (Figure 6D, Movie 4).

These results indicate that budding off and release of Piwi from mitotic nuage occurs along the central spindle MTs. We hypothesized that the interaction of nuage with the central spindle MTs facilitates Piwi’s dissociation from nuage. To more directly test this idea that MTs mediate dissociation of Piwi from mitotic nuage, we sought to allow cells to exit mitosis in the absence of MTs. To achieve this, we combined colcemid treatment with a *mad2* mutantion. Colcemid treatment effectively depolymerize MTs, which would normally cause mitotic arrest due to a spindle assembly checkpoint (Li et al., 2010). However, when this is combined with a *mad2* mutantion, cells exit mitosis even in the absence of MTs (Figure 6 – figure supplement 1). Under this condition, we found that Piwi remained in cytoplasmic nuage after mitotic exit, demonstrating that Piwi’s release from mitotic nuage and return to the nucleus is mediated by MTs (Figure 6E, F).

### *klp10A* is required for depolymerization of central spindle MTs

The above results led us to hypothesize that central spindle serves as a platform of Piwi dissociation from mitotic nuage such that Piwi can return to the nucleus. How does Klp10A, which is a kinesin motor that bends and depolymerizes MTs (Rogers et al., 2004), regulate this process? Because central spindle MTs seemed to facilitate dissociation of Piwi from mitotic nuage (Figure 6), and because Klp10A localizes to central spindle (Figure 4E), we investigated whether Klp10A may regulate the integrity of the central spindle.

To test this idea, we examined the morphology of the central spindle in control vs. *klp10A^RNAi^* testes. First, we found that the frequency of GSCs or SGs that contain central spindle MTs was significantly higher in *klp10A^RNAi^* compared to control testes (Figure 7A-C), suggesting that Klp10A promotes depolymerization of central spindle MTs. In control germ cells, disassembly of the central spindle and completion of S phase, as assessed by pulse labeling with 5-ethynyl-2'-deoxyuridine (EdU), are nearly all in synchrony. Most of the S phase cells (EdU+) contain central spindle (Figure 7D, arrowhead), whereas all post-S phase cells (EdU-) have resolved central spindle (Figure 7F). In a stark contrast, in *klp10A^RNAi^* testes, about half of post-S phase cells still contained the central spindle (Figure 7E, arrow, Figure 7F), indicating that central spindle disassembly is delayed in the absence of Klp10A. Importantly, increased frequency of germ cells with the central spindle is not due to skewed composition of cell cycle stages in *klp10A^RNAi^* germ cells, because *klp10A^RNAi^* germ cells had comparable frequency of being in S phase (22.6±8.6% EdU^+^ in control, *n*=89 GSC vs. 24.6±15.6% EdU+ in *klp10A^RNAi^*, n=103 GSC, n.s.) or in M phase (0.22±0.03 mitotic GSCs/testis in control, n=160 vs. 0.194±0.01 mitotic GSCs/testis in *klp10A^RNAi^*, n=222, n.s.).

**Figure 7.**
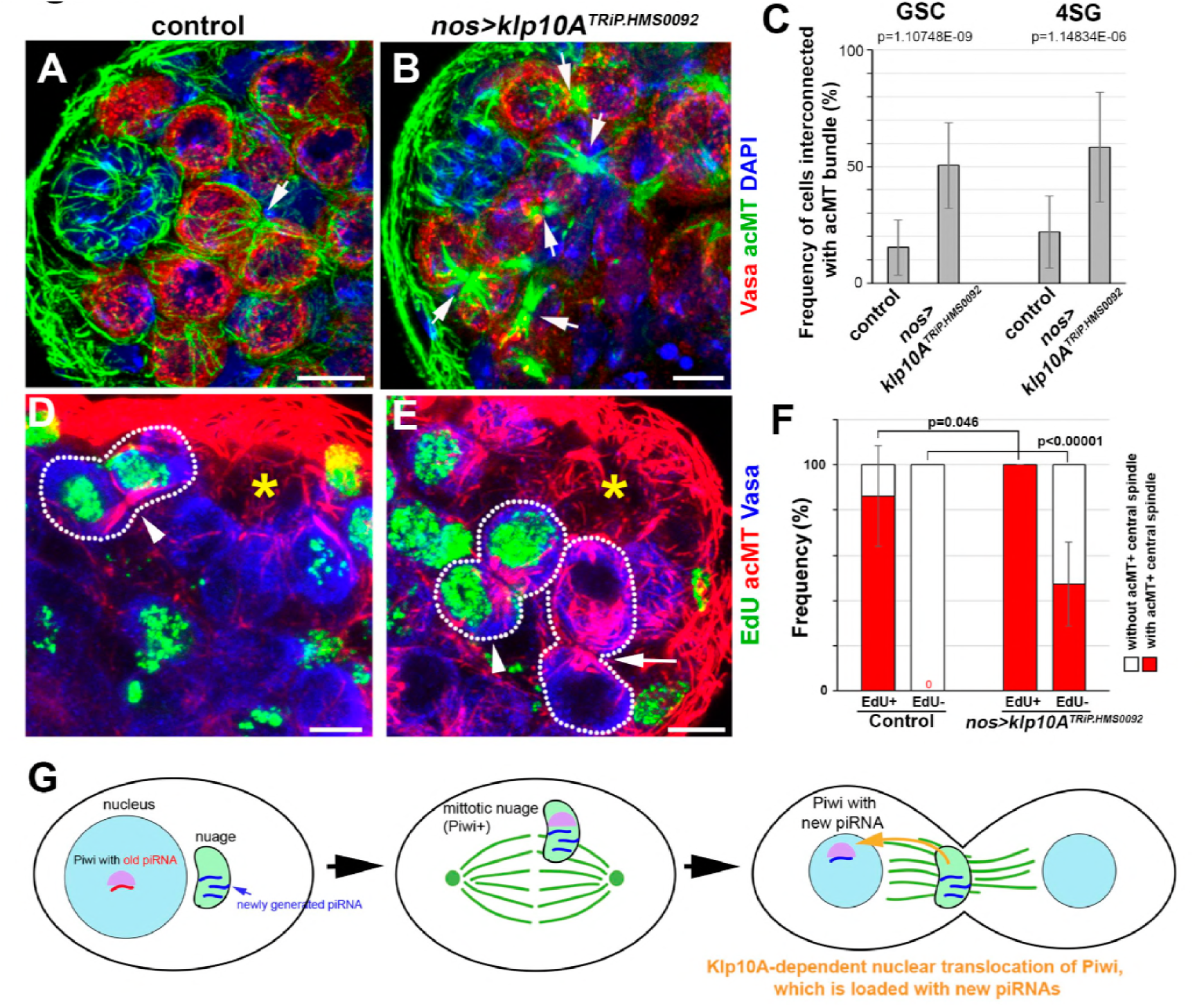
Klp10A promotes depolymerization of central spindle MTs. A, B) Apical tip region of wild type (A) and *klp10A^RNAi^* (B) testes, stained for Vasa (red), acMTs (green) and DAPI (blue). Arrows indicate bundle of acMTs between interconnected daughter cells. Bars: 5 *μ*m C) Frequency of acMT bundles in GSCs (n=181 for control, n=201 for *klp10A^RNAi^*) or 4-cell SGs (n=128 and n=95). Error bars indicate SD. p values of t-tests are provided. D-E) Apical tip region of a wild type (D) and a *klp10A^RNAi^* testis (E) with EdU (green) incorporation after 45min incubation period. Testes were co-stained for acMTs (red) and Vasa (blue). Arrowheads indicate EdU-positive GSC-GB pairs with acMT bundle. Arrow indicates EdU-negative GSC-GB pairs with acMT bundle in *klp10A^RNAi^*. Asterisks indicate the hub, dotted lines indicate GSC-GB pairs. Bars: 5 *μ*m. F) Frequency of acMT bundle in EdU-positive (n=32 for control, n=24 for *klp10A^RNAi^*) and EdU-negative GSCs (n=69 for control, n=79 for *klp10A^RNAi^*). Error bars indicate SD. p values of t-tests are provided. (G) model of how cell-cycle-dependent compositional changes of nuage is facilitated in SGs in a Klp10A-dependent manner.

Taken together, these results show that *klp10A* is required for depolymerization of central spindle MTs. Based on the observation that Piwi dissociates from mitotic nuage along the central spindle as it disassembles, we speculate that Piwi’s dissociation from central spindle is driven by MT depolymerization, which is facilitated by Klp10A, a MT-depolymerizing kinesin. Understanding how depolymerizing MTs can facilitate dissociation of Piwi from mitotic nuage awaits future investigation.

## Discussion

In this study, we revealed unexpected dynamic localization of piRNA pathway components during cell cycle of mitotically proliferating germ cells (GSCs and SGs) in the *Drosophila* testis. Piwi is nuclear in interphase, but associates with nuage specifically during mitosis, then returns to the nucleus at the end of the mitosis. We propose that Piwi’s interaction with nuage during mitosis might facilitate the loading of Piwi with piRNAs. Additionally, by interacting with nuage at every mitosis period, Piwi may gain an access to ‘updated’ repertoire of piRNAs that reflect currently active TEs (Figure 7G).

It is of note that this mechanism appears to be distinct from the cell biological mechanism of Piwi loading with piRNAs in the nurse cells of the female germ line (Malone and Hannon, 2009). Nurse cells are highly polyploid post-mitotic cells and thus do not undergo nuclear envelope breakdown as do SGs (Spradling, 1993). Piwi needs to be loaded with piRNAs to translocate into the nucleus (Handler et al., 2011; Olivieri et al., 2010; Wang et al., 2015). However, Piwi’s localization to nuage is not visible in wild type cells, likely because Piwi only transiently associates with nuage prior to nuclear translocation. Only when the piRNA pathway is completely compromised, as in the *aub ago3* double mutant, can we detect the failure of Piwi to enter the nucleus and appear accumulated in nuage (Wang et al., 2015). In contrast to this mechanism of piRNA loading and nuclear translocation of Piwi in post-mitotic nurse cells, our finding suggests that mitotically-dividing germ cells (SGs) in the testis may utilize mitosis (nuclear envelope breakdown) as a means to load Piwi with piRNAs. Nuclear envelope breakdown may represent a robust and efficient way of loading Piwi with piRNAs in SGs that divide every 12 hours.

Our data suggest that the ‘nuage cycle’ in the mitotic germ cells of the testis is partly facilitated by disassembly of central spindle MTs: Klp10A regulates depolymerization of the central spindle MTs during telophase, and Piwi dissociates from nuage at the depolymerizing central spindle MTs. How can depolymerizing MTs facilitate dissociation of Piwi from the nuage compartment? Recently, it was shown that liquid droplets of Tau protein concentrate tubulin dimers to facilitate MT formation, and that MT itself within the Tau droplets exhibit liquid-like properties (Hernández-Vega et al., 2017). Given that nuage is also a phase-separated compartment (Brangwynne et al., 2009; Nott et al., 2015), it is tempting to speculate that tubulin (generated from depolymerization of MTs by Klp10A) might influence the characteristics of nuage, lowering its affinity with Piwi. In addition, MTs have been shown to regulate the assembly, maturation, and disassembly of stress granules, another phase-separated, non-membrane bound compartment similar to nuage, although the underlying molecular mechanisms remain elusive (Chernov et al., 2009; Ivanov et al., 2003; Loschi et al., 2009; Shao et al., 2017). Taken together, we propose that Klp10A-dependent MT depolymerization at the central spindle facilitates Piwi’s dissociation from nuage to promote its translocation back into the nucleus. In summary, the present study reveals a novel cell biological mechanism by which nuclear and cytoplasmic arms of piRNA biogenesis communicate during the cell cycle, possibly allowing the coordination of transcriptional and post-transcriptional repression of target transcripts.

## Acknowledgements

We thank Elizabeth Gavis, Katalin Fejes Tóth, Dorothea Godt, Mikiko Siomi, the Bloomington Stock Center and the Developmental Studies Hybridoma Bank for reagents, the Yamashita lab members, Rebecca Tay, Margaret Starostik, Sue Hammoud and Lei Lei for the discussion and comments on the manuscript. This research was supported by Howard Hughes Medical Institute (to Y.Y. and S.E.J.) and NIH R01GM118875 (to J.K.K.).

## Materials and methods

### Fly husbandry and transgenic flies

Flies were raised in standard Bloomington medium at 25°C. The following stocks were used. *UAS-gfp-klp10A* (Inaba et al., 2015a), *vas-gfp* (Sano et al., 2002), *piwi-gfp* (a gift from Katalin Fejes Tóth)(Le Thomas et al., 2013), *mCherry-vas* (a gift from Elizabeth Gavis)(Lerit and Gavis, 2011), and the following stocks, obtained from the Bloomington Stock Center*: nos-gal4* (Van Doren et al., 1998), *UAS-upd* (Zeidler et al., 1999), *UAS-gfp* (Spana E., 1999, personal communication to FlyBase, ID: FBrf0111645), *UAS-aub-gfp* (Harris and Macdonald, 2001), *UAS-mCherry-*α-*tubulin* (Rusan and Peifer, 2007), *UAS-aub^TRiP.GL00076^*, *UAS-klp10A^TRiP.HMS00920^* (Flybase: FBrf0214641, FBrf0212437), *mad2^EY21687^* (Li et al., 2010), Df(3L)BSC437 (FlyBase: FBrf0204472)

### Immunoprecipitation and western blotting

For immunoprecipitation, 150 pairs of testes were lysed in 0.5 ml HEPES based buffer [50mM HEPES pH 7.5, 250mM NaCl, 0.1% Nonidet P40, 0.2% Triton X-100, supplied with cOmplete protease inhibitor mixture (Roche)]. In Klp10A co-immunoprecipitation (co-IP) experiments, the cleared lysates were incubated with rabbit anti-Klp10A serum (1:200, (Chen et al., 2016)) for 4 hours, extended by 2 extra hours after supplement with Protein A Dynabeads (Life Technologies). The beads were washed four times with lysis buffer and proteins were eluted with SDS-PAGE protein sample buffer. For co-IP experiments with GFP tagged proteins, the cleared lysates were incubated with GFP-Trap magnetic beads (Chromotek) according to manufacturer's instructions. To detect precipitated proteins on western-blots, the following primary antibodies were combined with horseradish peroxidase conjugated secondary antibodies (Abcam): guinea-pig anti-Klp10A (1:10,000, (Chen et al., 2016)), anti-GFP (1:3000, ab290, Abcam,), anti-Vasa d-26 (1:3000, Santa Cruz Biotechnology).

### Mass-spectrometry

To purify Klp10A-interacting proteins from early germ cells, we used testes expressing GFP-Klp10A and Upd to enrich GSCs/early germ cells (*nos-gal4>UAS-upd, UAS-gfp-klp10A*) (Kiger et al., 2001), and GFP-Klp10A was purified using anti-GFP nanobodies, followed by liquid chromatography-tandem mass spectrometry (LC-MS/MS) (Neumuller et al., 2012). 1500 pairs of 0-2 days old *nos-gal4*>*UAS-upd, UAS-gfp-klp10A* or *nos-gal4*>*UAS-upd, UAS*-*gfp* (control) flies were used. Dissected testes were stored in Schneider’s Insect Medium (Sigma-Aldrich) at ambient temperature up to two hours. All the downstream steps were performed at 4°C, unless otherwise noted. Collected testes were homogenized with Dounce homogenizer in 1 ml lysis puffer [10mM Tris-HCl pH 7.5; 150mM NaCl; 0.5 mM EDTA; 0.5% Nonidet P40, supplied with 1 mM PMSF (Sigma) and cOmplete protease inhibitor cocktail (Roche)]. Lysate was incubated for 30 minutes, and cleared by two times 10 min-centrifugation at 16.000x g. Supernatant were stored at −80 °C at this stage. After thawing on wet ice, 1.5 ml dilution buffer (10mM Tris/Cl pH 7.5; 150 mM NaCl; 0.5mM EDTA) was added to the supernatant prior to adding 25 μl packed GFP-Trap agarose beads (Chromotek). After 3 hours of incubation, beads were washed three times with dilution buffer and proteins were eluted with 2x sample buffer. The eluate was boiled for 5 min and subjected to SDS-PAGE. The entire volume of the eluate was separated in a single lane of an 8-16% Tris-Glycine SDS-PAGE gradient minigel (Life technologies). After electrophoresis the unstained lane has been cut in two at around 70kDa and the gel parts were washed for three times 15min in ddH20. Proteins were digested over-night with 5 ng/μL Trypsin (Promega) in 50 mM ammonium bicarbonate at 37 °C. The peptides were extracted using 5% v/v formic acid and acetonitrile subsequently. The extracts were dried in a speed vacuum and cleaned using ZipTip C18 (Millipore). LC-MS/MS analysis was conducted by a LTQ Orbitrap XL mass spectrometer (Thermo Scientific) at Fred Hutchinson Cancer Research Center, Proteomics Resource. Data were analyzed with Proteome Discoverer 1.4 (Thermo Scientific), with the peptide spectral matches filtered to 5% false discovery rate (FDR).

### Immunofluorescence staining and microscopy

Fixation and immunofluorescence staining of testes was performed as described previously (Chen et al., 2016). The following primary antibodies were used. Mouse anti-Fasciclin III (FasIII; 1:100; 7G10, developed by Goodman C, obtained from Developmental Studies Hybridoma Bank; (Patel et al., 1987)), rabbit anti-Vasa (1:200; d-26, Santa Cruz Biotechnology), mouse anti-acetylated-Tubulin (1:100, 6-11B-1, Sigma), rabbit anti-phosphorylated (Thr3) histone H3 (PH3) (1:200; clone JY325, Upstate), mouse anti-LaminB (1:200; C20; Santa Cruz Biotechnology), guinea pig anti-Traffic jam (Tj; 1:400; a gift from Dorothea Godt; (Li et al., 2003)), rabbit and guinea-pig anti-Klp10A (1:3000/1:1000 respectively; (Chen et al., 2016)). Alexa Fluor-conjugated secondary antibodies were used (1:200; Thermo Fisher Scientific), EdU was detected by Click-iT Plus EdU Imaging Kit with Alexa Fluor 647 (Thermofisher). Testes were mounted into VECTASHIELD media with 4',6-diamidino-2-phenylindole (DAPI; Vector Labs). Images were captured using a Leica TCS SP8 confocal microscope with a 63×oil-immersion objective (NA=1.4) and processed by ImageJ software (Schindelin et al., 2012).

### RNA in situ hybridization

RNA Fluorescent *in situ* Hybridization (RNA FISH) on testes was performed to detect *HOBO* transcripts by applying Stellaris DNA oligo probe set conjugated with Quasar 570 fluorophore (Biosearch Technologies). Probe sequences are listed in Supplementary Table S1. RNA FISH was performed by applying a method described by (Raj and Tyagi, 2010) on the *Drosophila* testes.

### *Ex vivo* treatment of *Drosophila* testis

To enrich cells in metaphase testes were dissected and transferred to Schneider's insect medium (Sigma) containing MG132 (20 μM final concentration, Sigma-Aldrich) (Moutinho-Pereira et al., 2010) or colcemid (100μM final concentration, Calbiochem) (Venkei and Yamashita, 2015). At this concentration of MG132 up to 4.5h incubation period, all types of mitotically active cells in the testes gradually accumulated in metaphase with intact bipolar spindle. With colcemid treatment, mitotically active cells were similarly arrested in mitosis without spindle due to MT depolymerization. For EdU incorporation, the dissected testes were incubated for 45min (10 μM, Thermofisher) in Schneider's insect medium (Gibco) at 25°C.

### RNA extraction, library preparations, and sequencing

Total RNAs were extracted by the Trizol method. 300 pairs of testes were used for each sample, and biological triplicates for each genotype was used. For small RNA sequencing, 2.5 μg of total RNAs were treated with DNase I (Amplification Grade, ThermoFisher) and recovered by RNA Clean & Concentrator-5 columns (Zymo). Libraries were generated from DNaseI treated RNAs using reagents from the TruSeq Small RNA Library Prep Kit (Illumina), while following a previously published protocol from (Wickersheim and Blumenstiel, 2013) that uses a terminator oligo to deplete *Drosophila* 2S rRNA from the final sequencing libraries. The libraries were sequenced in one lane on a HiSeq 2500 machine (Illumina) at the UCLA Broad Stem Cell Research Center BioSequencing Core. The terminator oligo complimentary to Drosophila 2S rRNA was ordered from IDT and the sequence is TAC AAC CCT CAA CCA TAT GTA GTC CAA GCA /3SpC3/.

For mRNA sequencing, 2.5 μg of total RNAs were treated with DNase I (Amplification Grade, ThermoFisher) and recovered by RNA Clean & Concentrator-25 columns (Zymo). DNaseI treated RNAs were then treated with Ribo-Zero Gold rRNA Removal Kit (Illumina). Libraries were generated from rRNA depleted RNAs using TruSeq Stranded Total RNA Sample Prep Kit (Illumina). The libraries were sequenced by a HiSeq 4000 machine (Illumina) at the UCLA Broad Stem Cell Research Center BioSequencing Core.

### Small RNA-seq data analysis

Three biological replicates of each condition were used to test for statistical significance and comparative analysis. Raw qseq files were converted to fastq format using custom Python and awk scripts. Small RNAs were clipped from 3’ adapter sequences using Trimmomatic 0.27 (Bolger et al., 2014). Reads mapping with at most 1 mismatch to rRNA and miRNA hairpin sequences were parsed out by Bowtie2 v2.2.3 (Langmead and Salzberg, 2012). The remaining reads were aligned to the *D. melanogaster* transcriptome (Dm3) with at most 1 mismatch allowed using Bowtie2 v2.2.3. Abundance estimation was done at annotated genes, piRNA clusters, and TEs using the eXpress pipeline within the piPipes suite (Han et al., 2015; Roberts and Pachter, 2012). We used Flybase for annotation of protein coding genes and TEs. piRNA cluster annotations were done according to (Brennecke et al., 2007). For piRNA analysis, we selected reads that were 23-29 nucleotides in length to ensure other endo-siRNA species did not influence downstream analysis. miRNA hairpin reads were used to normalize between libraries. Differential expression analysis was done in DESeq2 3.7 using the Wald test and adjusted p-values were corrected using the Benjamin-Hochberg procedure with an FDR threshold of 0.01(Love et al., 2014). Plots were generated using R. Sequencing data is available at GEO: accession number: GSE122596.

### Ping-pong activity analysis

piRNA reads were mapped directly to the *D. melanogaster* transcriptome (Dm3) using Bowtie2 v2.2.3 (Langmead and Salzberg, 2012). BAM files from Bowtie2 were converted to BED files using several pipelines, including BEDTools 2.7 from the piPipes suite, which assigns nucleotide positions of each read across TE transcript bodies (Han et al., 2015; Quinlan and Hall, 2010). In addition to nucleotide position, reads were categorized as sense and antisense. Sense was defined as piRNAs that were derived from cleavage of the TE mRNAs and antisense was defined as piRNAs that were antisense to annotated TE transcripts. Custom Python scripts were used to calculate the number of 5’ to 5’ complementarity between sense and antisense reads. Ping-pong ratios were calculated for each transposon feature across all libraries. To calculate the ping-pong ratio of each transposon, we used custom Python scripts to calculate the number of piRNAs in which sense piRNAs with an A at the 10-nt position or antisense piRNAs with a U at the 1-nt position showed 10 nt of complementarity from the 5’ end. We then divided the number of such pairs by the total number of piRNA reads. The resulting ratio allowed for quantification of ping-pong activity, without needing to normalize for library size. The calculation of ping-pong ratio was done using custom Python scripts. Plots were generated using R.

### mRNA-seq data analysis

Raw qseq files were converted to fastq format using custom Python and awk scripts. Due to the repetitive nature of transposon sequences, reads were aligned to the *D. melanogaster* genome (Dm3) by STAR-2.6.0a while allowing for up to 100 multi-alignments per read (Dobin et al., 2013). Feature and abundance estimation were determined using TEtranscripts 2.0.3 (Jin et al., 2015). TEtranscripts initially distributes multi-mapped reads evenly among potential matches and optimizes the distribution of multi-mapped reads using the Expectation Maximization approach. Variance was measured based on normalized gene expression counts and we performed principal components analysis and found samples derived from wild-type clustered separately from *Klp10A-RNAi* on Principle Component 1 (Figure 2 - Figure Supplement 2A). Differential analysis was done using DESeq2 3.7 using the Wald test and p-values were corrected using the Benjamin-Hochberg procedure with an FDR threshold of 0.01 (Love et al., 2014). All Plots were generated using R. Pipeline of Analysis is described in (Figure 2 - Figure Supplement 3A and 3B).

### Live imaging

Testes from newly enclosed flies were dissected in Schneider’s Drosophila medium (Gibco) and prepared for live imaging as described previously (Cheng and Hunt, 2009). The testis tips were placed into a drop of medium in a glass-bottom chamber and were covered by regenerated cellulose membrane (Spectrum Lab). The chamber was mounted on a three-axis computer-controlled piezoelectric stage. An inverted Leica TCS SP8 confocal microscope with a 63× oil immersion objective (NA = 1.4) was used for imaging. Live imaging was performed at ambient room temperature. Images were processed using ImageJ software.

**Figure 2 – Figure Supplement 1.**
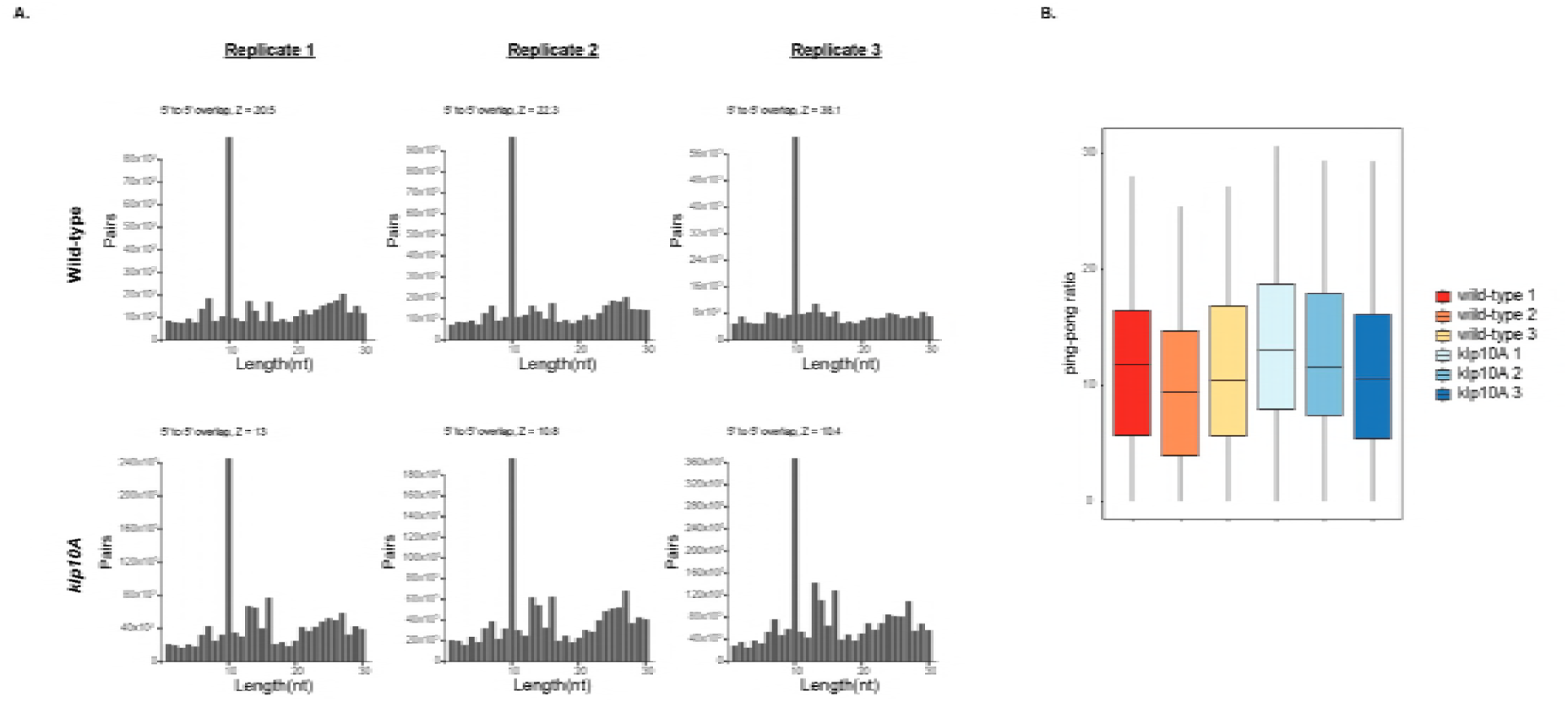
Ping-pong pathway activity remain unaltered after *klp10A* depletion. A) Histogram showing the distribution of antisense and sense piRNA pairs of piRNAs mapping to transposons. B) The box-plots are showing the distribution of ping-pong ratios of each transposon. Each box-plot is a different biological replicate. The Ping-pong ratio of each transposon was calculated by taking the sum of piRNA reads in which sense piRNAs with a 10 nt A and antisense piRNAs with a 1nt U showing 10 nucleotide compementarity from the 5’ end and dividing it with the total number of piRNA reads.

**Figure 2 – Figure Supplement 2.**
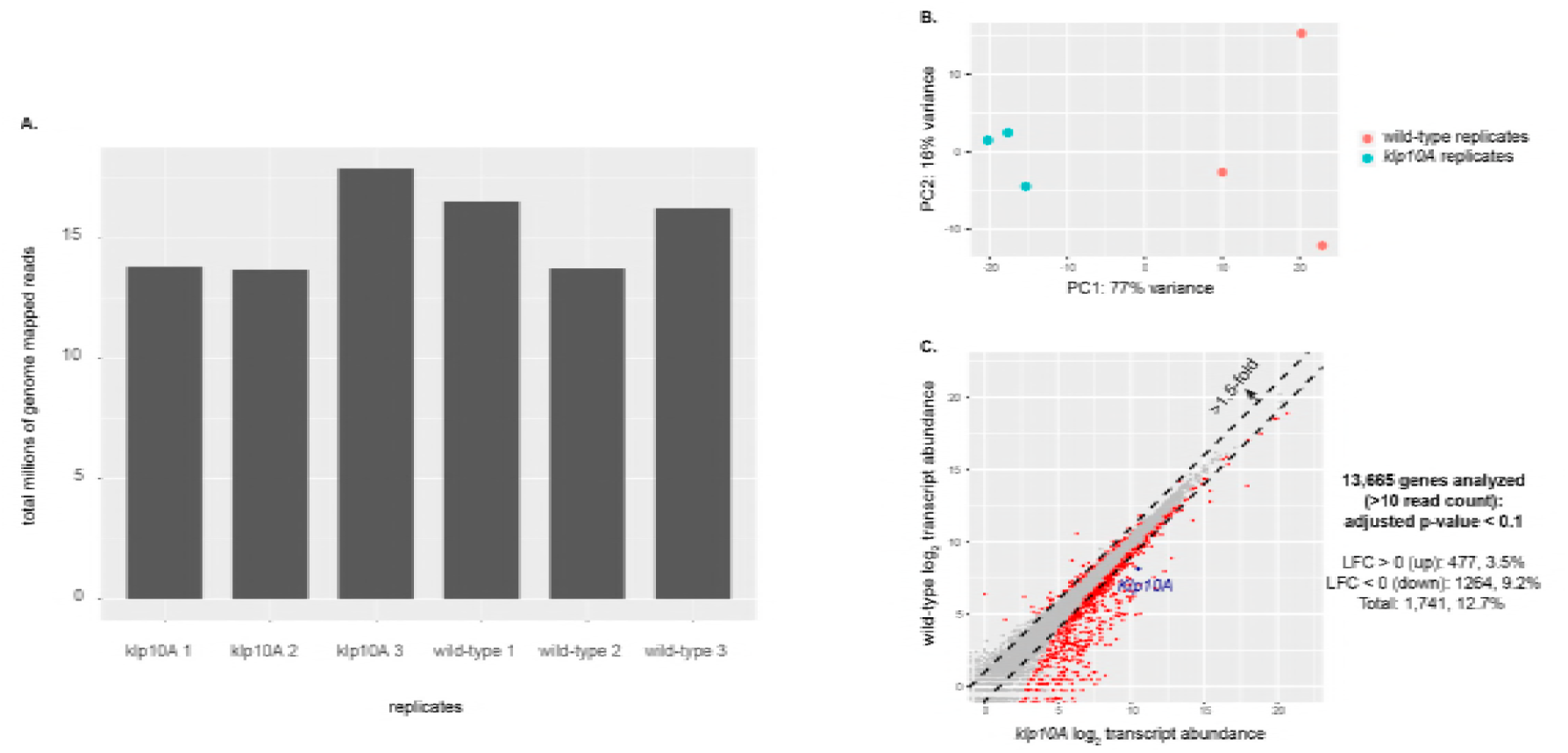
Characterization of RNAseq datasets. A) Bar plot showing number of total genome mapped reads per RNAseq library B) Clustering analysis of wildtype (n=3 replicates) and *klp10A* (n=3 replicates) RNAseq libraries. PC, principal component; POV, percentage of variance. C) Scatter plot showing mean genic read abundance of *klp10A^RNAi^* versus wild-type libraries. LFC: log2 fold change.

**Figure 2 – Figure Supplement 3.**
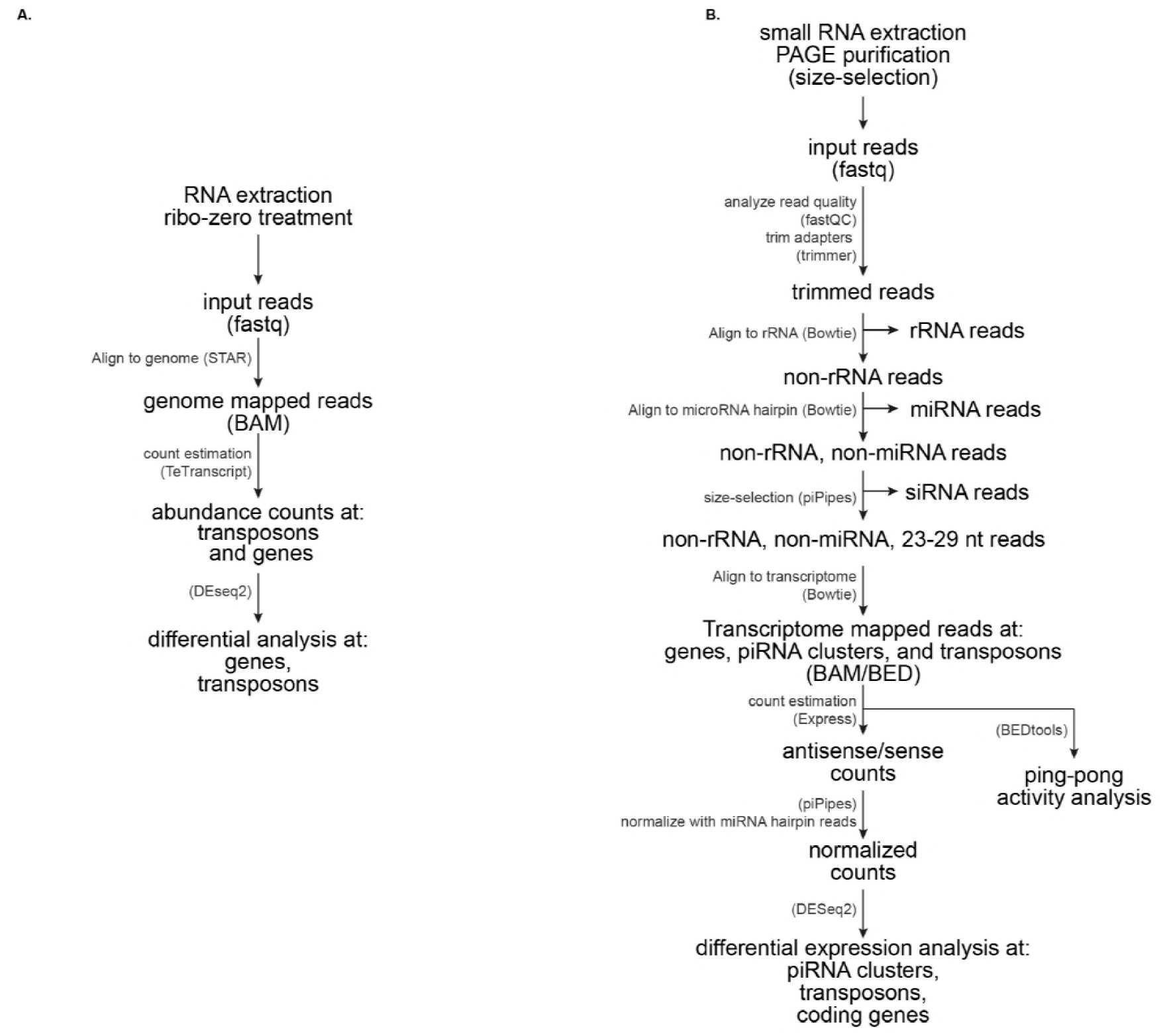
Pipeline of small RNA and RNA sequencing analysis. A) Pipeline outlining RNA sequencing analysis. B) Pipeline outlining small RNA sequencing analysis.

**Figure 3 – Figure Supplement 1.**
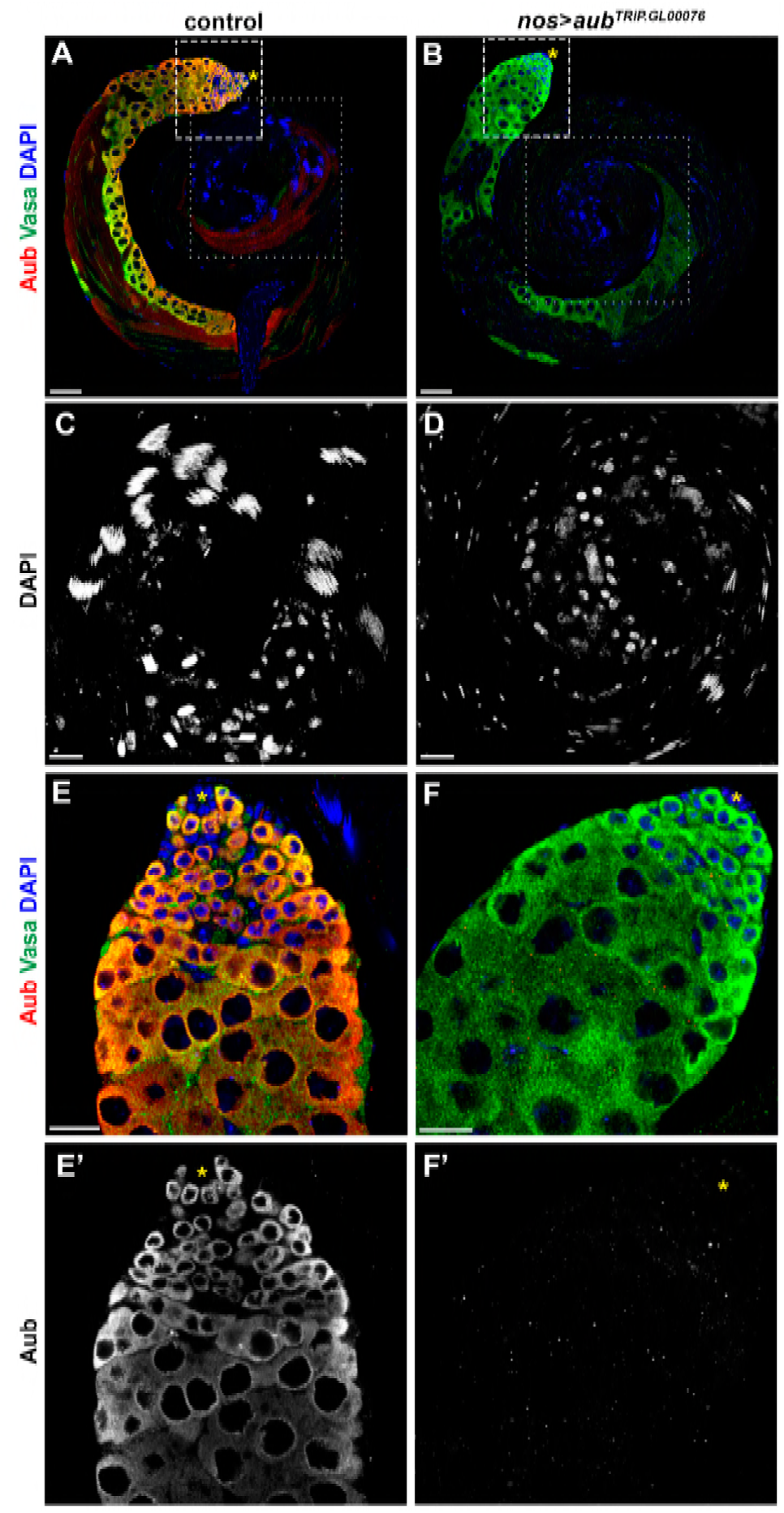
Knockdown efficiency of *aub* RNAi. In wild type testis *aub* expression is restricted to the germ line (A, E and E’; ref1). *nos*-*gal4* driven *aub^RNAi^* depletes Aub in all germ cells (B, F and F’). Knock down of *aub* results in sperm head clustering defect, which is characteristic for piRNA pathway mutant testes. Aub (red), Vasa (green), DAPI (blue). C and D zoom in the dotted line marked regions of A and B. E-F’ zoom in the broken line indicated regions of A and B. A and B bars: 50 μm, C-F: 20 μm.

**Figure 3 – Figure Supplement 2.**
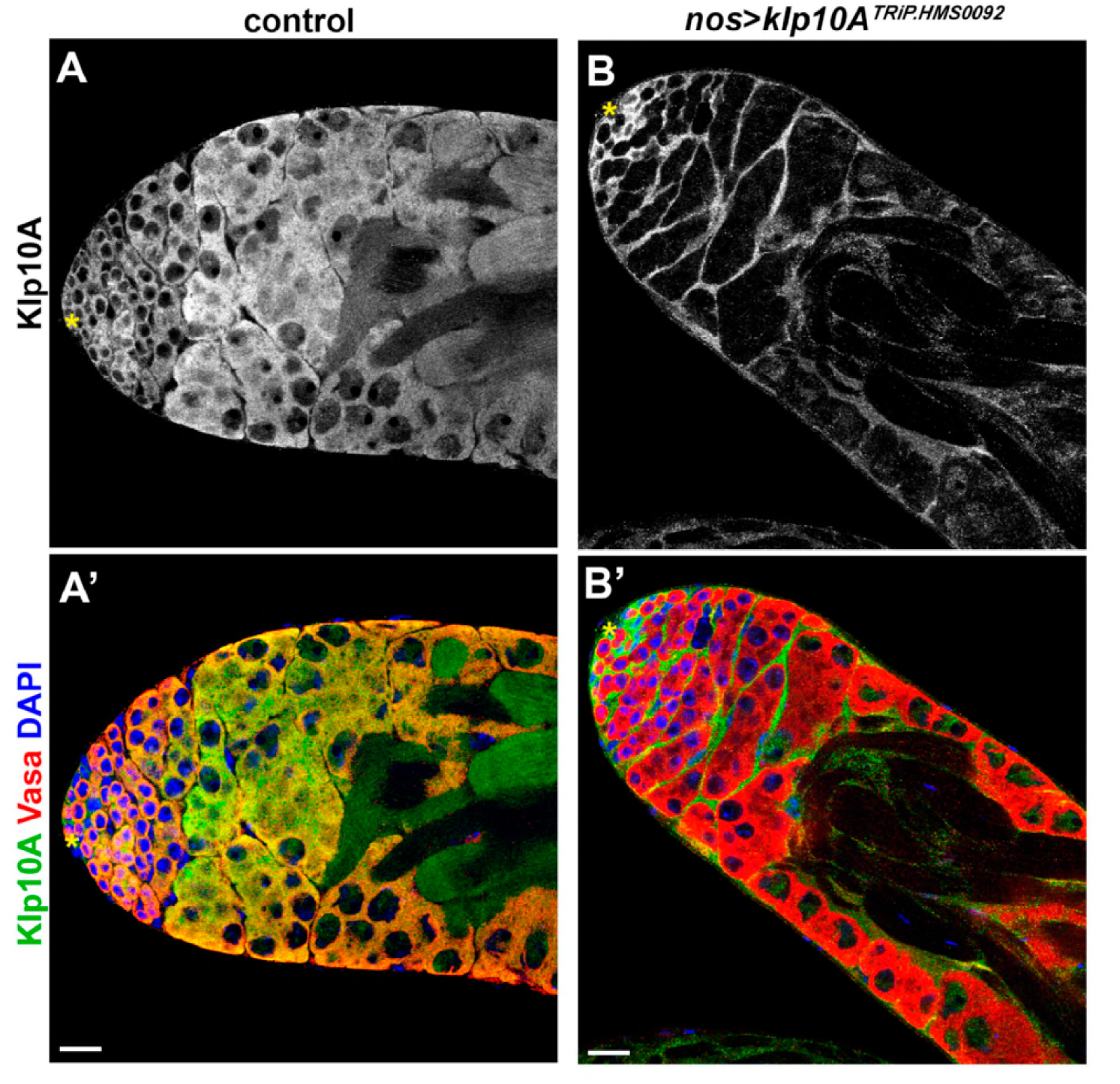
Knockdown efficiency of Klp10A by RNAi. In wild type testis Klp10A is abundant in the apical tip region (A-A`). *nos*-*gal4* driven *klp10A^RNAi^* depletes Klp10A in all germ cells within the testis (B-B`). Note that Klp10A signal in the somatic cells remain. Vasa (red), Klp10A (green), DAPI (blue). Bars 20 μm.

**Figure 4 – Figure Supplement 1.**
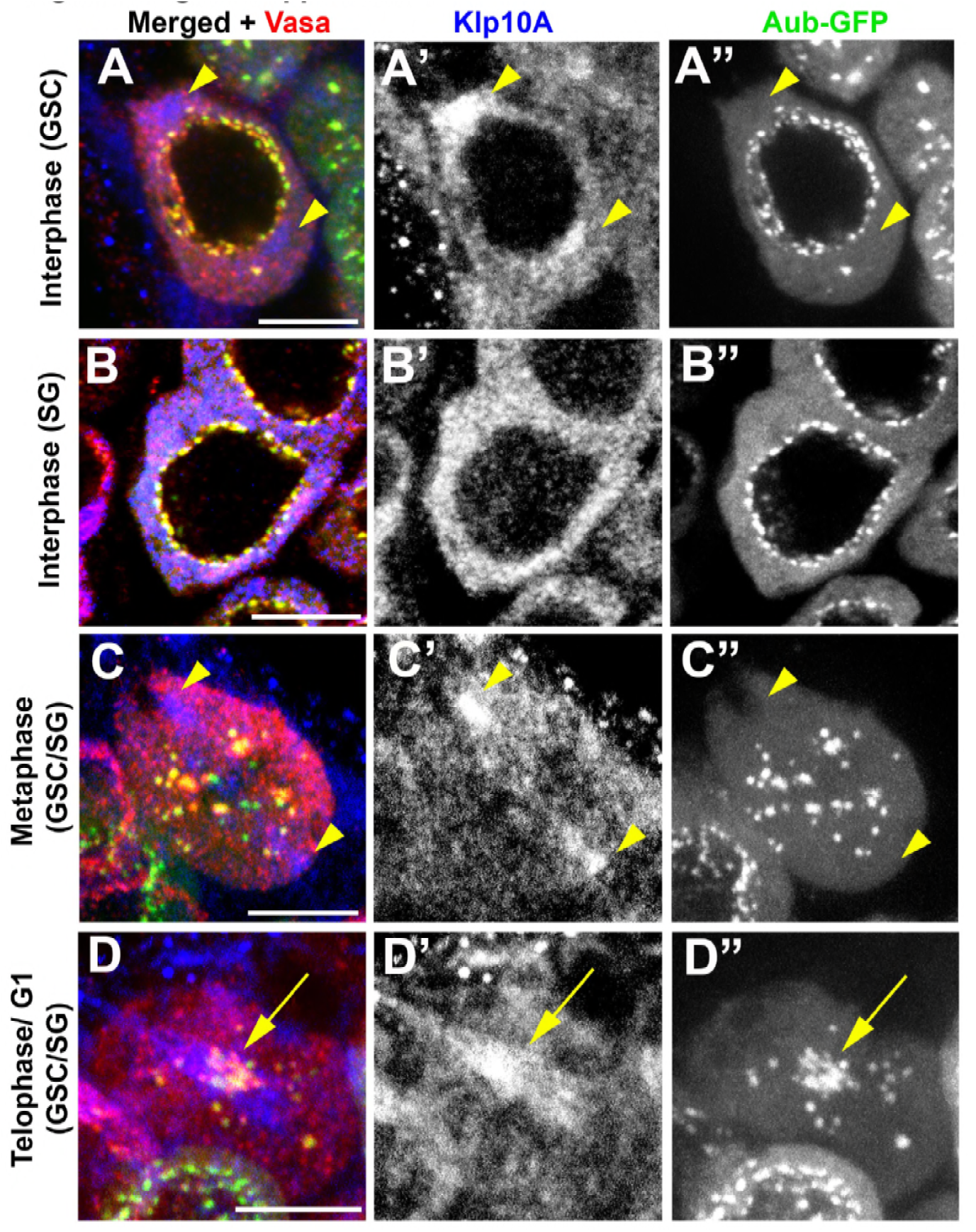
Colocalization of Klp10A and piRNA pathway components Vasa and Aubergine during the cell cycle. Klp10A (blue), Vasa (red) and Aub-GFP (green) in the germline. Interphase GSC (A-A”), Interphase SG (B-B”), GSC/SG in metaphase (C-C”), and GSC/SG in telophase/G1(D-D”). Klp10A localization to centrosomes is indicated by arrowheads and central spindle localization with arrows. Bars: 5 *μ*m.

**Figure 5 – Figure Supplement 1.**
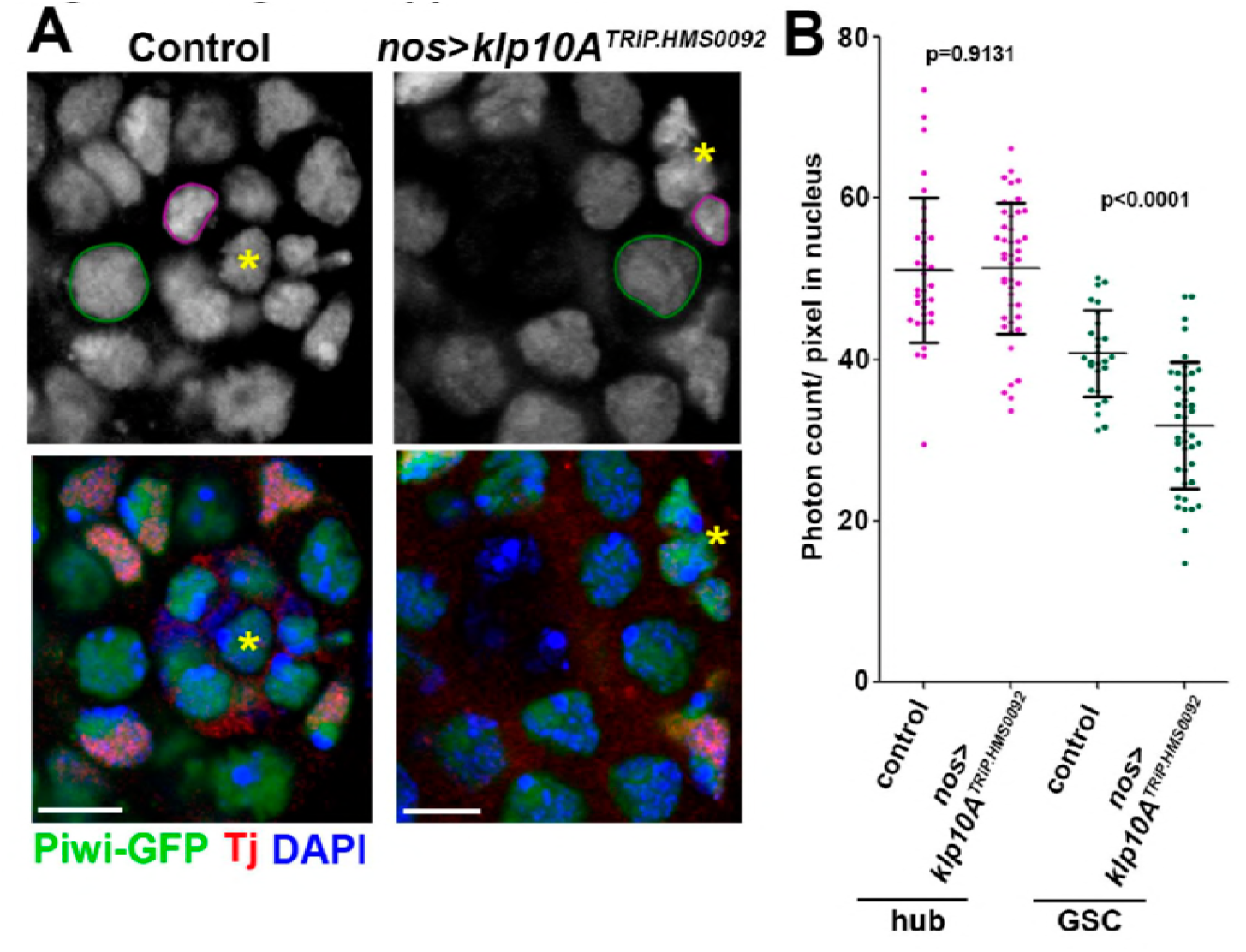
Nuclear Piwi level is decreased upon knockdown of *klp10A.* A) Piwi-GFP (green) in the apical tip of control vs. *klp10A^RNAi^* testes. Tj (red) identifes cyst stem cells and early cyst cells), DAPI (blue). Green lines encircle GSC nuclei. Magenta lines encircle hub cell nuclei. Bars: 5 μm. Asterisks indicate hub. B) Photon counts in nuclear areas of hub cells (magenta) and GSCs (green) in control vs. *klp10A^RNAi^*. Error bars indicate SD, p values from t-tests are provided. As *klp10A* is knocked down only in germline (with *nos-gal4*), Piwi nuclear level is expected to be unchanged in hub cells, which were confirmed by photon counting, and this serves as an internal control. On the contrary, Piwi nuclear level was reduced in GSCs upon knockdown of *klp10A*. Note that similar reduction in the nuclear Piwi was observed in SGs as well: however, we used hub cells and GSCs to quantify Piwi levels, because their juxtaposition allows the most accurate comparison between two cells.

**Figure 6 – Figure Supplement 1.**
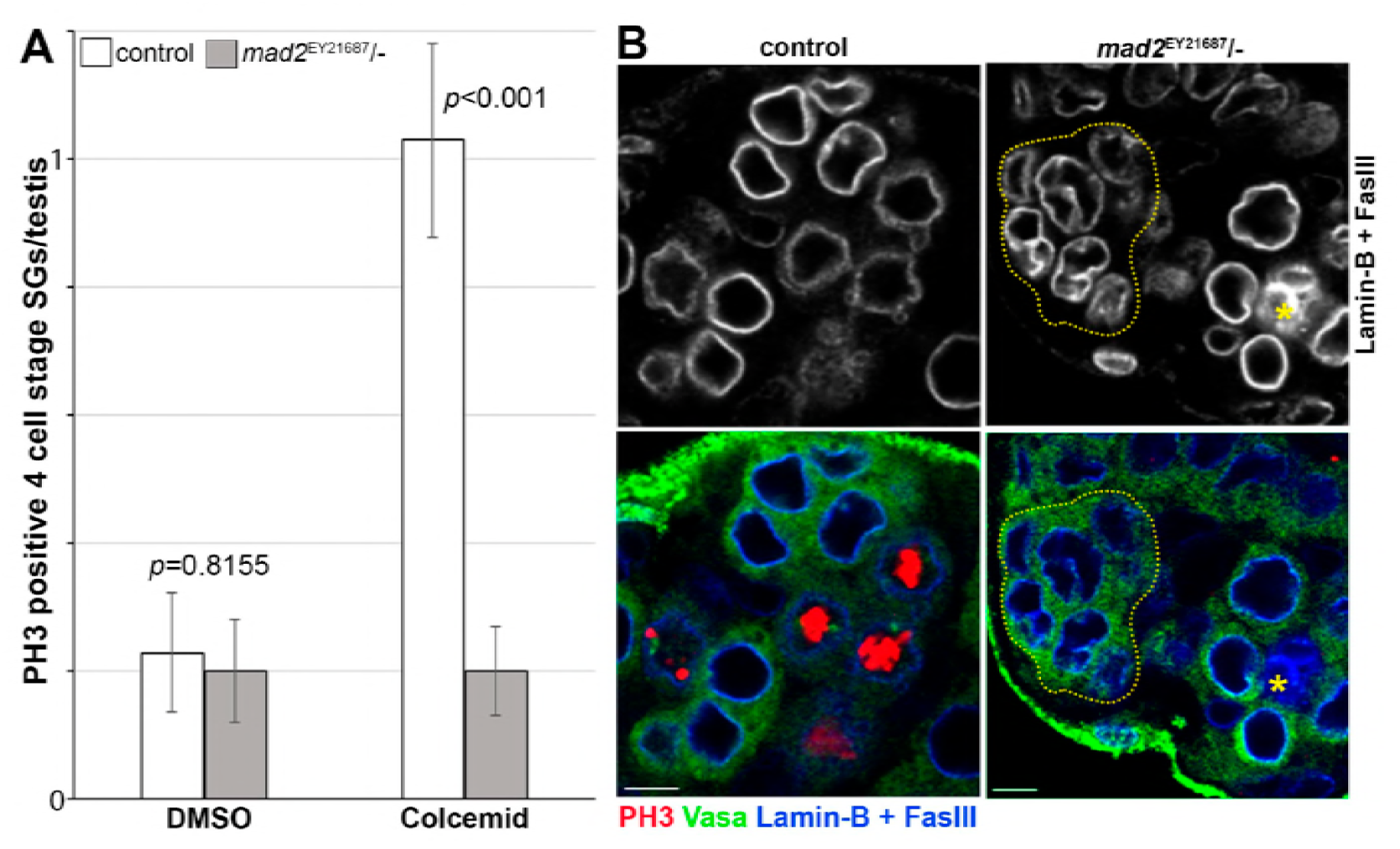
SGs exit mitosis in the absence of MTs after the elimination of the spindle assembly checkpoint. A) *mad2* mutant SGs do not arrest in mitosis after MT depolymerisation. B) After prolonged MT depolymerisation some SGs own enlarged, presumably polyploid, post-mitotic nuclei in *mad2* mutants. One SG with enlarged nuclei is encircled by dotted line. Vasa (green), PH3 (red), Lamin-B and FasIII (blue). Bars: 5 μm.

**Movie 1. Dynamics and composition of mitotic nuage in wild type SGs.**

Dynamics of Piwi-GFP (green) and mCherry-Vasa (red) in mitosis of wild type SGs. Time 0 is set to anaphase B onset. Z-stacks were collected with the indicated time intervals with Z-step size 0.5 *μ*m and 9 steps per time point. Bar on first frame: 5 *μ*m. Note that at the end of mitosis, cytoplasmic nuage is Vasa+, Piwi-.

**Movie 2. Dynamics and composition of mitotic nuage in a GSC an SG cell upon knockdown of *klp10A***.

Dynamics of Piwi-GFP (green) and mCherry-Vasa (red) in mitosis of *klp10A*^RNAi^ GSC (left bottom) and SG (upper right). Time 0 is set to anaphase B onset. Z-stacks were collected with the indicated time intervals with Z-step size 0.5 μm and 11 steps per time point. Bar on first frame: 5 *μ*m. Note that at the end of mitosis, cytoplasmic nuage is Vasa+, Piwi+, although some portion of Piwi returns to the nucleus.

**Movie 3. Piwi disassembly along the central spindle MTs in a wild type GSC during mitotic exit**.

Lime-lapse live imaging of Piwi-GFP (green) and mCherry-Tubulin (red) in a wild type GSC. Time 0 is set to first frame (0 min) in anaphase of the cell cylce. Z-stacks were collected with the indicated time intervals with Z-step size 0.5 μm and 8-12 steps per time point (note that when Piwi+ somatic nuclei move into the focal plane, those frames were manually removed). Bar on first frame: 5 *μ*m.

**Movie 4. Defective Piwi disassembly in telophase SGs upon knockdown of *klp10A*.**

Lime-lapse live imaging of Piwi-GFP (green) and mCherry-Tubulin (red) in *klp10A*^RNAi^ GSCs. Time 0 is set to first frame (0 min) in telophase/G1 of the cell cycle. Z-stacks were collected with the indicated time intervals with Z-step size 0.5 μm and 6-10 steps per time point (note that when Piwi+ somatic nuclei move into the focal plane, those frames were manually removed). Bar on first frame: 5 *μ*m.

**Supplementary Table 1.** Stellaris DNA oligo probes for RNA in situ.

Source Data

Figure 1-Source Data 1: Mass-spectrometry results of GFP pull down from *nos*>*upd gfp* (control) and *nos*>*upd gfp*-*klp10A* flies.

